# Alternative polyadenylation is a determinant of oncogenic Ras function

**DOI:** 10.1101/2020.06.08.140145

**Authors:** Aishwarya Subramanian, Mathew Hall, Huayun Hou, Marat Mufteev, Bin Yu, Kyoko E. Yuki, Haruka Nishimura, Anson Sathaseevan, Benjamin Lant, Beibei Zhai, James Ellis, Michael D. Wilson, Mads Daugaard, W. Brent Derry

**Affiliations:** Developmental and Stem Cell Biology Program, Peter Gilgan Centre for Research and Learning, The Hospital for Sick Children, Toronto, M5G 0A4, ON, Canada; Genetics and Genome Biology Program, Peter Gilgan Centre for Research and Learning, The Hospital for Sick Children, Toronto, M5G 0A4, ON, Canada; Department of Molecular Genetics, University of Toronto, Toronto, M5S 1A8, ON, Canada; Department of Medicine, University of Toronto, Toronto, M5S 1A8, ON, Canada; Vancouver Prostate Centre, Vancouver, V6H 3Z6, BC, Canada; Department of Urologic Sciences, University of British Columbia, Vancouver, V5Z 1M9, BC, Canada

## Abstract

Alternative polyadenylation of pre-mRNA has been recently shown to play important roles in development and cancer. Activating mutations in the Ras oncogene are common drivers of many human cancers but the mechanisms by which they cooperate with alternative polyadenylation are not known. By exploiting the genetics of *C. elegans*, we identified *cfim-1/CFIm25*, a subunit of the alternative polyadenylation machine, as a key determinant of hyperactive Ras function. Ablation of *cfim-1* increased penetrance of multivulva phenotype in *let-60/Ras* gain-of-function (gf) mutant through shortening of transcripts at the 3’ untranslated region, including p21 activated kinase *pak-1/PAK1* and multidrug transporter *mrp-5/ABCC1*. Depletion of CFIm25 in human KRAS-driven cancer cells resulted in a similar shortening of 3’ untranslated regions in the *PAK1* and *ABCC1* transcripts, which caused an epithelial-to-mesenchymal transition and increased cell migration. Exploiting the mechanisms by which alternative polyadenylation affects activated oncogene output could offer novel approaches for the treatment of Ras-driven tumors.

## INTRODUCTION

The transition of normal cells to the neoplastic state requires genetic alterations that cause activation of oncogenes and loss of tumor suppressors. Genome-wide analysis of numerous cancers has yielded a dizzying array of secondary mutations in genes that can impact the evolution of the cancer cell and its tumorigenic potential^1,2^. The Ras oncogene is conserved from yeast to human and is one of the most frequently mutated genes in cancer, with up to 30% of all tumors possessing gain-of-function mutations in one or more of the three Ras paralogs: *KRAS, HRAS and NRAS*^3–5^. Ras proteins are critical regulators of signaling pathways that control proliferation, cell survival, and tissue differentiation. Point mutations in the GTPase domain of Ras lead to constitutive activation of pro-survival pathways that contribute to tumor development and progression^4,6^. Ras and other oncogenes, such as MYC and Notch, require specific levels of expression, pathway activation, or cross-talk with other signaling axes to sustain tumor phenotypes in mammalian systems^7,8^. Studying the mechanisms by which these signaling pathways interact is challenging using established cell lines, which are often highly mutated and aneuploid^9,10^. Model organisms provide robust complimentary systems to study genetic interactions within the context of living animals.

The powerful genetic tools of the nematode worm *Caenorhabditis elegans* have provided much insight into the *in vivo* functions of oncogenes such as Ras^11^. Mutational activation of the *C. elegans* Ras homolog *let-60* occurs by a glycine to glutamic acid substitution at amino acid 13 (the n1046 allele), similar to the glycine to valine at amino acid 12 found in numerous human cancers^12^. The hallmark phenotype of this *let-60* gain-of-function (gf) allele is the multivulva (Muv) phenotype, in which ectopic vulvae develop along the ventral body of the worm^13^. Forward genetic screens in *C. elegans* have helped establish the hierarchical order of Ras/MAPK signaling components, and uncovered numerous genes that regulate Ras/LET-60 signaling that are conserved from worm to human^14–16^.

Although substantial efforts to target and directly inhibit Ras in cancer have been undertaken, these ventures have shown limited success^17,18,19^. Likewise, strategies for targeting downstream Ras effectors have had limited clinical success due to dose-limiting toxicity of the compounds^20–22^. In addition, genetic compensation and signaling crosstalk within the Ras signaling network contributes to drug resistance in many tumors^23,24,25^. As such, novel strategies to inhibit Ras signaling for cancer treatment are needed to overcome these obstacles. While ambitious genome sequencing efforts have uncovered variants in genes that can impact the tumorigenic state, much less is known about how epigenetic influences on gene expression and function cooperate with oncogenic pathways.

An important epigenetic mechanism of gene regulation involves an essential step in mRNA maturation, which determines the length of the 3’ untranslated region (3’UTR). This occurs by endonucleolytic cleavage of the pre-mRNA in the 3’UTR, followed by addition of the poly(A) tail. This cleavage reaction is directed by poly(A) site (PAS) consensus sequences composed of multiple *cis* elements, including *i)* an AAUAAA hexamer recognized by the cleavage and polyadenylation specificity factor (CPSF) complex; *ii)* downstream U/GU-rich elements that are recognized by the cleavage stimulation factor (CstF); and *iii)* upstream U-rich elements recognized by cleavage factor Im (CFIm).These components establish a platform for subsequent recruitment of cleavage factor IIm (CFIIm) and its co-factors to form the functional 3’UTR processing holoenzyme that cleaves and polyadenylates pre-mRNA^26^. Many genes contain multiple polyadenylation sites in their 3’UTRs, enabling cleavage and polyadenylation at different PAS locations, a process termed alternative polyadenylation (APA). APA serves to direct the placement of the poly(A) tail such that isoforms of the same gene are produced with different 3’UTR lengths. The 3’UTR contains numerous regulatory sites for the binding of microRNAs and RNA binding proteins (RBPs), which regulate mRNA stability, localization, and translational efficiency^27,28^.

Shorter 3’UTRs of specific genes have been associated with higher rates of proliferation, and these transcripts undergo 3’UTR lengthening as the cells reach a more differentiated state^29^. About 30-40% of genes in *C. elegans* and about 70% of genes in mammals undergo APA, respectively^30,31^. Alterations in APA have also been associated with tumorigenesis, where transformed cells tend to have shorter 3’UTRs of key proliferative and survival genes. This often results in higher translation rates and increased levels of proteins that promote tumorigenesis^32^. Apart from regulating transcript stability and translation, recent studies have demonstrated how changes in APA of specific transcripts can also influence the localization and functions of resulting proteins^33,34^. These observations have provided insight into the complexity of APA in diseases such as cancer; however, the precise mechanisms by which APA factors cooperate with activated oncogenes remains unknown.

In this study, we uncover evolutionarily conserved roles of the APA factor, *cfim-1/CFIm25* in oncogenic Ras signaling using *C. elegans* and human cancer cell models. We showed that *cfim-1/CFIm25* functions to buffer oncogenic Ras output by regulating the ratio of 3’UTR lengths for conserved transcripts in both systems. These include the p21 activated kinase (*pak-1/PAK1*) and the multi-drug resistance protein (*mrp-5/ABCC1*), where shortening of their 3’UTRs cooperates with oncogenic Ras to enhance its functional output in development and tumorigenic cell behaviour.

## RESULTS

### RNAi screen uncovers *cfim-1* as a novel regulator of activated *let-60*/Ras

The multivulva (Muv) phenotype used to measure hyperactive Ras activity in *C. elegans* was quantified in two ways: (i) by quantifying the percentage of worms with at least one ectopic vulva (% Muv), and (ii) by generating a vulva induction score^35^. Vulva induction was measured by counting the number of cells that have adopted vulval fates at the fourth larval (L4) stage (described in Methods). In normal development only 3 vulval precursor cells (VPCs) adopt vulval cell fates, thus the induction score for wild-type worms is 3.00 and the % Muv is 0 (**Fig. 1a, b**). Under conditions of hyperactive LET-60/Ras, more VPCs adopted vulval cell fates, so induction scores above 3 represent Muv worms. *let-60(gf) (n1046)* worms that possessed strong hyperactive Ras function in the somatic tissue had an average vulval induction score of 4.09 and were 71% Muv (**Fig 1a, b, c**).

**Fig. 1.**
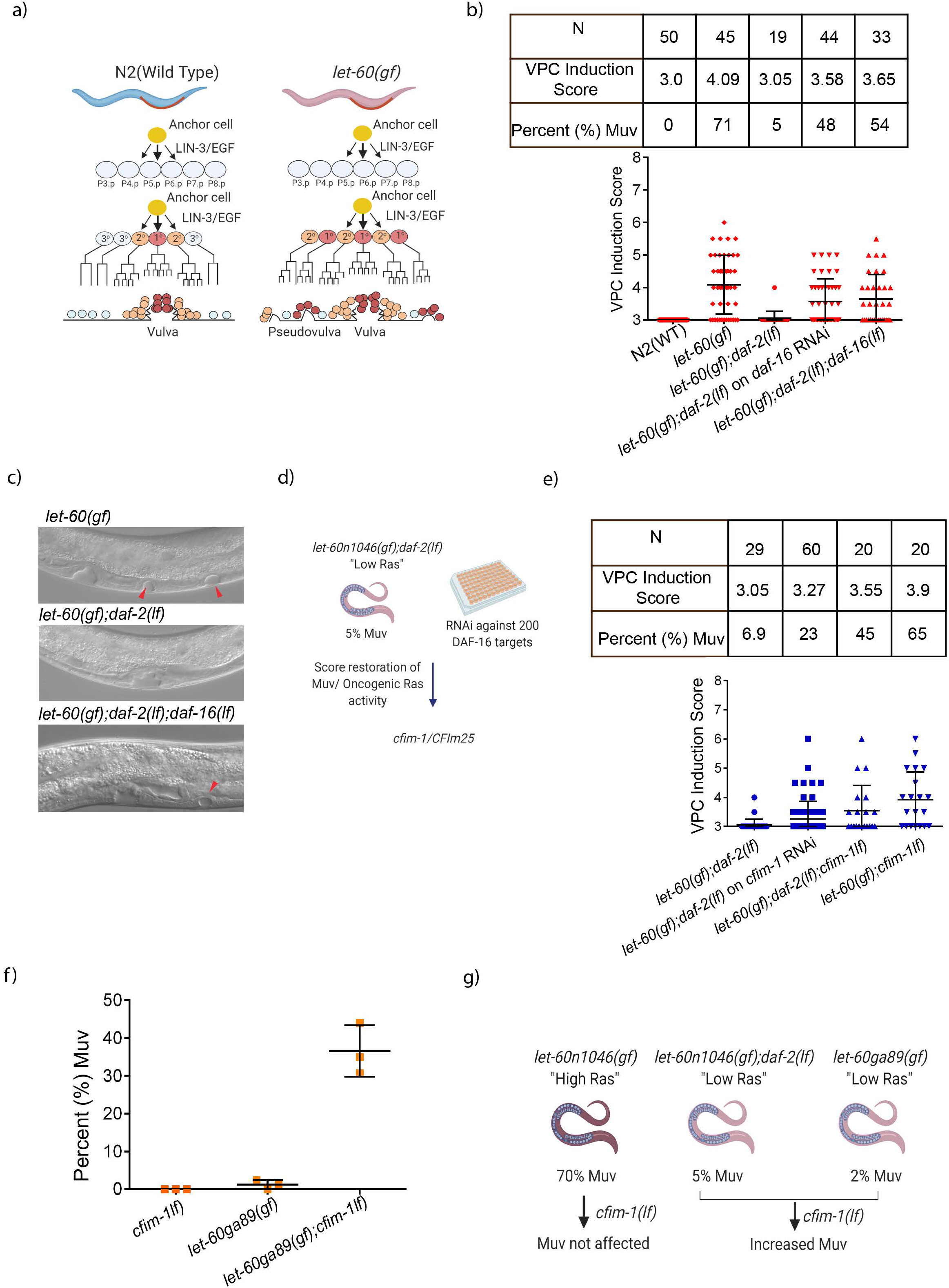
*cfim-1* regulates oncogenic Ras activity in *C. elegans*. a) Schematic outline of vulval development in *C.elegans* b) VPC induction scores for N2 and combinations of *let-60(gf), daf-2(lf),* and *daf-16(lf)* mutants. % Muv indicates percentage of animals with more than 3 VPCs induced. N indicated the number of animals measured. Y-axis origin is set to 3 to represent the normal number of VPCs that are induced to divide. c) Representative images of Larval 4 stage worms expressing the hyperactive *let-60(gf)* allele, *let-60(gf)*;*daf-2(lf)* double mutant in which the Muv phenotype was completely suppressed and *let-60(gf); daf-2(lf); daf-16(null)* triple mutant in which the Muv phenotype was partially restored. Red arrows denote ectopic pseudo-vulvae. d) Schematic representation of screen performed by feeding *let-60(gf);daf-2(lf)* worms (supressed MUV) RNAi against DAF-16 targets e) VPC induction scores for *let-60(gf);daf-2(lf)* on HT115 control and *cfim-1* RNAi. RNAi against *cfim-1* showed partial restoration of MUV and a stronger effect was observed in *a let-60(gf);daf-2(lf);cfim-1(lf)* strain. *cfim-1(lf)* in the context of *let-60n1046(gf)* showed no effect. % Muv indicates percentage of animals with more than 3 VPCs induced. N indicates the number of animals measured. Y-axis origin is set to 3 to represent the normal number of VPCs that are induced to divide. f) Percentage (%) Muv scored for the *let-60ga89(gf)* and *let-60ga89(gf); cfim-1(lf)* mutants. Each data point represents quantification from three independent experiments g) Schematic representation of the effect of *cfim-1* loss in the context of different levels of Ras activation

We previously demonstrated that the hypersensitivity to DNA damage-induced germline apoptosis of the Ras/*let-60(ga89)* allele, which has a more penetrant phenotype in the germline, is dependent on a functional IGF1R homologue *daf-2.* Loss-of-function (lf) mutation in *daf-2* completely suppresses sensitivity to DNA damage-induced germline apoptosis in *let-60(ga89)* mutants^36^. This dependency on *daf-2/IGF1R* was also observed in the soma, where *daf-2(lf)* suppressed the Muv phenotype of *let-60(n1046gf)* worms to 7% (**Fig. 1b, c**)^37^. This suppression was partially dependent on the transcription factor *daf-16/FOXO3a*, which functions downstream of the *daf-2/IGF1R* pathway. *daf-2(lf); let-60(gf); daf-16(null)* triple mutants had a restoration of the Muv phenotype to a 3.65 induction score and 54% Muv (**Fig 1b, c**). Taking advantage of the *let-60(gf); daf-2(lf)* double mutant in which the Muv phenotype was suppressed, we conducted a screen to identify novel genes that restore the Muv phenotype; similar to what was observed by ablating *daf-16* (**Fig. 1d**).

We focused on genes that are either known or predicted to be DAF-16 transcriptional targets, reasoning that restoring the Muv phenotype in *let-60(gf)*; *daf-2(lf)* mutants would identify transcriptional targets of DAF-16 that regulate LET-60/Ras function. Therefore, a list DAF-16 transcriptional target genes was generated from several unbiased studies, including Chromatin Immunoprecipitation (ChIP) and DNA Adenine Methyltransferase Identification (DamID). Genes in the top 5% of methylation peaks from the DamID study and at least one other study were considered potential DAF-16 targets^38,39,40–44^. Based on this, we screened 202 candidate genes by RNA interference (RNAi) and found the ablation of *cfim-1* increased the Muv phenotype of *let-60(gf)*; *daf-2(lf)* double mutants to an induction score of 3.27 and 23% Muv (**Fig. 1e**). This effect was even more pronounced after we generated a CRISPR deletion allele of *cfim-1* and built a *let-60(gf); daf-2(lf); cfim-1(lf)* triple mutant, which was 45% Muv and had an induction score of 3.55 (**Fig. 1e**). Interestingly, *cfim-1* loss in the context of the *let-60(n1046)* single mutant (high Ras activation) did not show any additional enhancement of Muv (**Fig. 1e**). Since the n1046 allele is a strong *let-60*/Ras(gf) mutant, we wondered if *cfim-1* might enhance a weaker *let-60* allele. Therefore, we crossed the *cfim-1(lf)* allele into the *let-60(ga89)* mutant and observed an increase in Muv from 2% to 40%, indicating that co-operation between *cfim-1* and *let-60* was independent of the *daf-2*/IGF1R pathway (**Fig. 1f**). Our data indicate that *cfim-1(lf)* can enhance the Muv output with low levels of oncogenic Ras/LET-60 in *C. elegans* (**Fig. 1g**).

### *cfim-1* regulates APA of transcripts that mediate oncogenic *let-60*/Ras output

*cfim-1* encodes the *C. elegans* homolog of mammalian *NUDT21/CFIm25*, a subunit of the 3’ polyadenylation cleavage CFIm complex that directs poly(A) tails to distal cleavage sites to produce transcripts with longer 3’UTRs. Depletion of *CFIm25* causes a global shortening in 3’UTRs, resulting in more stable transcripts that escape negative regulation by microRNAs and/or RBPs (**Fig. 2a**)^45^. Therefore, we next asked if *cfim-1* modulates *let-60*/Ras signaling output through biased APA of specific transcripts.

**Fig. 2.**
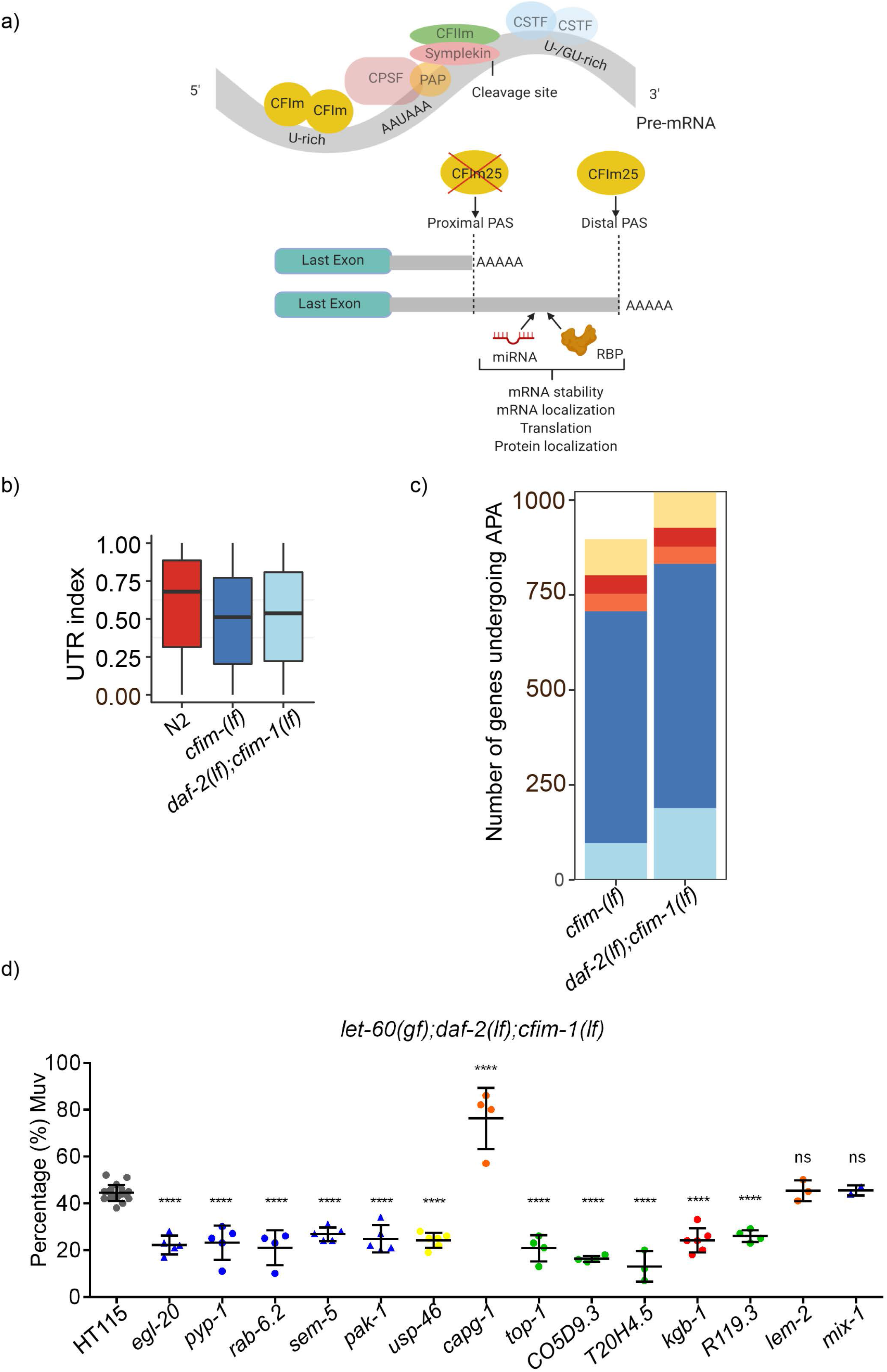
*cfim-1* loss of function promotes 3’UTR shortening of let-60/Ras pathway-related transcripts. a) Schematic outline of CFIM-1/CFIm25 function in the context of APA. CFIm25 is a component of the CFIm complex (denoted in yellow) which together with other protein complexes dictates polyadenylation of transcripts. CFIm25 biases towards the usage of distal polyadenylation sites, resulting in a longer 3’UTR. The longer 3’UTR often contains regulatory sequences such as miRNA and RNA binding protein sites which regulates transcript stability and translation. Loss of CFIm25 is associated with shortening of 3’UTRS and subsequent loss of regulatory sequences. b) 3’UTR index (proportion of reads mapping to the most distal polyadenylation site (PAS)) calculated in *cfim-1(lf)* and *daf-2(lf); cfim-1(lf)* strains. UTR indexes close to one represent longer 3’UTRs. P value < 2.2e-16 compared to N2 control. Data is the average from three independent replicates of each strain. c) DexSeq analysis and quantification of number of genes altered by APA. “Shorter” and “longer” denote increased expression of this particular isoform in the respective strains as described in Supplementary Fig. 2. d) RNAi screen of the Ras pathway-associated genes in the *let-60(gf);daf-2(lf);cfim-1(lf)* mutant. Each data point represents an independent experiment. Colours/shapes of data points correlate with indicated colours/shapes of genes (representing functional categories) in network analysis (**Supplementary Fig. 2**).

We reasoned that if *cfim-1* functions to attenuate *let-60*/Ras signaling output, changes in the 3’UTRs of genes in *cfim-1(lf)* mutants would uncover transcripts encoding proteins that either feed back to the Ras/MAPK pathway or modulate downstream signaling cascades. Worms were subjected to whole genome 3’UTR sequencing and downstream analysis was performed by calculating the UTR index (mapping reads to the most distal PA site). Our analysis revealed that the *cfim-1(lf)* single and *daf-2(lf); cfim-1(lf)* double mutants exhibited significant overall 3’UTR shortening, where a significant portion of genes had lower number of reads mapping to the most distal PA site (**Fig. 2b, Supplementary Fig. 1b**). In contrast, the *daf-2(lf)* and *daf-2(lf); daf-16(lf)* mutants did not exhibit significant alterations in their 3’UTR landscapes compared with wild-type controls (**Supplementary Fig. 1a, b**). Given that *cfim-1* was initially screened as a predicted DAF-16 target, we investigated the transcript levels of *cfim-1* and discovered that it was not altered in *daf-2(lf)* and *daf-2(lf); daf-16(lf)* strains compared to the known DAF-16 targets, indicating that *cfim-1* was not likely a transcriptional target of DAF-16 (**Supplementary Fig. 1c**). This data, together with *cfim-1(lf)* restoring Muv in the *let-60(ga89)* strain, independent of *daf-2*/IGF1R, indicates that *cfim-1* functions downstream, or in a parallel pathway, to attenuate activated LET-60/Ras output.

We next asked how the altered 3’UTR landscape in the absence of *cfim-1* might modulate Muv in the *let-60(gf)* mutant. In addition to the UTR index, we also employed the DEXSeq method to classify APA events. Poly(A) sites within 3’UTR of a gene were denoted as “exons” and the number of reads mapping to these poly(A) sites was quantified. This enabled us to classify genes into different APA classes, such as isoform switching (increase in short and decrease in long, or vice versa), or changes in just one isoform (short/long), between *cfim-1(lf)* mutants compared to wild-type controls (**Supplementary Fig. 2**). Consistent with the UTR index result, loss of *cfim-1* significantly altered the APA landscape (**Fig. 2c)**. We identified 493 genes with shortened 3’UTRs in both *cfim-1(lf)* single and *daf-2(lf); cfim-1(lf)* double mutants (denoted as “shorter in mutants”), and 186 genes shortened specifically in the *daf-2(lf); cfim-1(lf)* double mutant (**Fig. 2c**). Gene Ontology analysis of these 679 genes revealed 97 that were enriched in Ras signaling or processes that regulate oncogenesis in mammalian cells, such as cell migration, cell cycle and cytoskeletal dynamics, as well as a number of genes involved in vulva development and kinase activity (**Supplementary Fig. 3**). We screened these 97 genes by RNAi in *let-60(gf)*; *daf-2(lf)*; *cfim-1(lf)* triple mutants and quantified the Muv phenotype. We uncovered 12 genes that modulated the Muv phenotype, of which 3 function in cell migration (*kgb-1/MAPK10*, *sem-5/GRB2* and *pak-1/PAK1*) and one in vulva development (*usp-46/USP46*). Interestingly, ablation of *capg-1*, a condensin complex subunit, increased the percentage of Muv (**Fig. 2e**). We also investigated these 12 genes in *let-60(n1046)* mutant and found no significant effect on the Muv phenotype, suggesting that these genes likely possessed *cfim-1(lf)* dependency (**Supplementary Fig. 4a**).

**Fig. 3.**
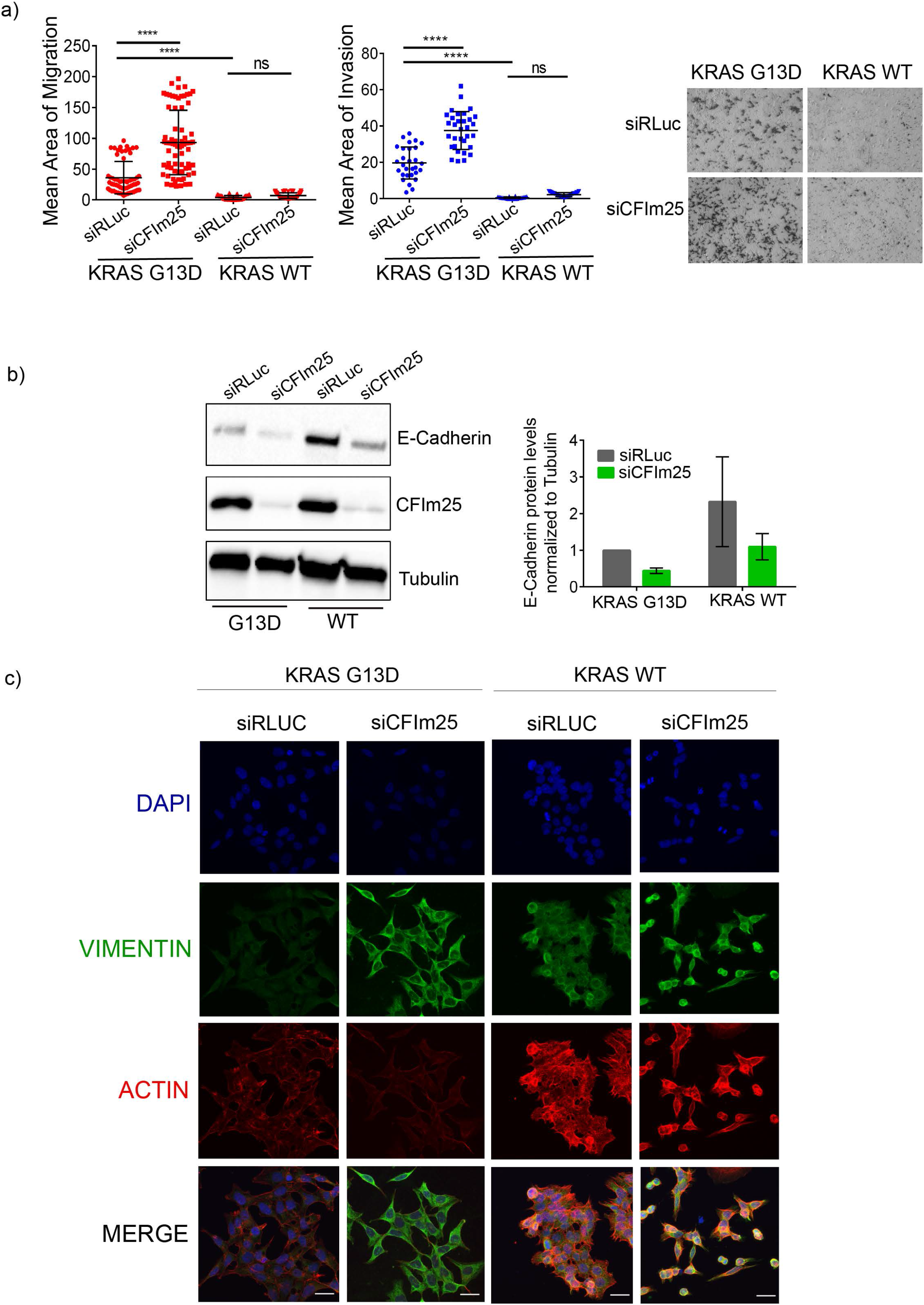
CFIm25 depletion increases migration and invasion of of KRAS mutant cells. a) Quantification of migration and invasion of KRAS G13D and KRAS WT HCT116 cell lines with/without CFIm25 knockdown. Each data point represents a field of view captured across three (migration) or two (invasion) independent experiments. ****p<0.0001, ns=not significant, One-way ANOVA with Tukey’s post hoc test. Images represent a single field of view of cell migration following knockdowns with indicated siRNAs. b) Representative western blot of E-cadherin expression in KRAS WT and KRAS G13D cell lines 72h post transfection with the indicated siRNAs and quantification of E-Cadherin protein levels normalized to Tubulin across two independent experiments. c) Representative Immunofluorescent images (maximum projection) of Vimentin (green) and actin (red) levels in KRAS G13D and KRAS WT 72h post transfection with the indicated siRNAs. Nuclei were stained with DAPI (blue). Scale bars represent 25μm

**Fig. 4.**
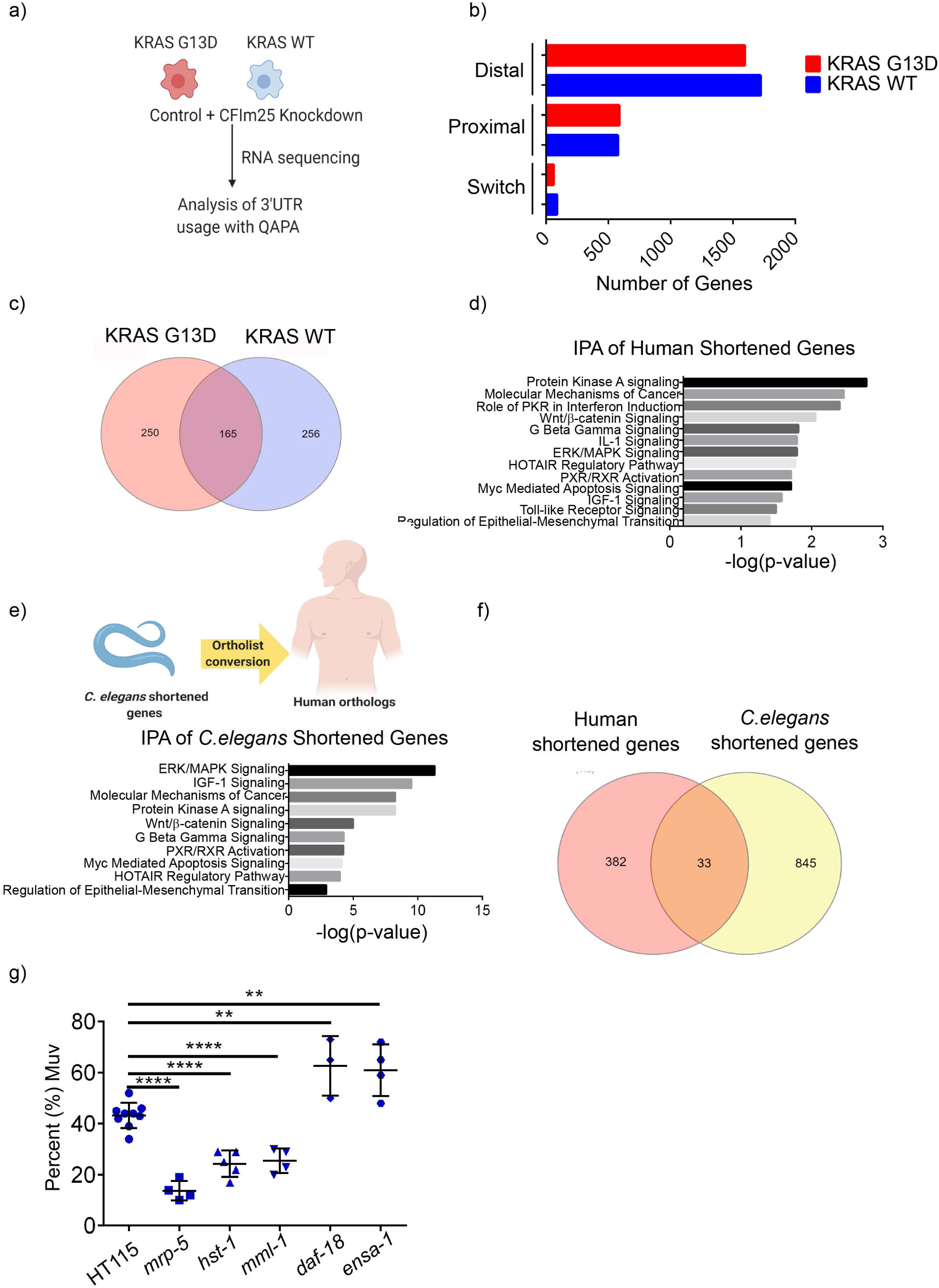
*cfim-1/CFIm25* regulates 3’UTR shortening of conserved genes in both *C.elegans* and human cancer. a) Schematic outline of RNA sequencing performed in KRAS G13D and KRAS WT HCT116 cells. Sequencing was performed with three independent replicates for each condition. b) Quantification of alterations in just one isoform (Distal or Proximal) or both isoforms (Switch) in KRAS G13D and KRAS WT cells. c) Venn diagram showing overlap in the number of genes undergoing 3’UTR shortening (increased expression of proximal isoform) in CFIm25-depleted KRAS G13D and KRAS WT cells. d) IPA analysis of the shortened transcripts in KRAS G13D cells. e) Schematic outline describing the conversion of *C. elegans* genes into their human orthologs, followed by IPA analysis. f) Overlay of *C.elegans* shortened genes with the human KRAS G13D shortened genes dataset to uncover commonly regulated genes. g) RNAi screen of commonly regulated *C.elegans* genes in the *let-60(gf);daf-2(lf);cfim-1(lf)* mutant. Each data point represents quantification from an independent experiment. ****p<0.001; **p<0.01, Unpaired Student’s t test

We further assessed the correlation between 3’UTR shortening and gene expression of the 12 *cfim-1*-dependent genes. Only *egl-20* showed increased gene expression with 3’UTR shortening in *daf-2(lf); cfim-1(lf)* double mutants, whereas the other 11 genes did not exhibit a significant correlation (**Supplementary Fig. 4b**). This suggests that 3’UTR shortening of those 12 genes could have other consequences, such as altered stability of specific isoforms, altered translation, localization and function^46^. Overall, our results indicate that *cfim-1* regulates APA of multiple transcripts that affect oncogenic *let-60/RAS* output in *C. elegan*s, and the human orthologs of these genes also have established roles in cancer (**Table 1**).

**Table 1.** Disease relevant *C. elegans* genes The functions of human orthologs of *C. elegans* genes that affect Muv phenotype downstream of *cfim-1*.

| ***C. elegans* Gene** | **Human Ortholog** | **Function** |
| --- | --- | --- |
| *egl-20* | *WNT16* | Cell fate regulator, paracrine signaling to promote tumor cell survival<sup>73,74</sup> |
| *top-1* | *TOP1, TOP1MT* | DNA topoisomerase, regulates DNA topology during transcription<sup>75</sup> |
| *C05D9.3* | *ITGB4* | Transmembrane glycoprotein receptors that regulate cell-cell adhesion, upregulated in many cancers<sup>76,77</sup> |
| *usp-46* | *USP12, USP46* | Deubiquitinating enzyme, possesses both oncogenic and tumor suppressor roles<sup>78,79</sup> |
| *kgb-1* | *MAPK10/JNK3* | Kinase involved in proliferation, differentiation and cell death, activated by inflammatory cytokines<sup>80</sup> |
| *T20H4.5* | *NDUFS8* | Complex I of respiratory chain, responsible for NADH oxidation<sup>81</sup> |
| *pyp-1* | *PPA1, PPA2* | Inorganic pyrophosphatase that plays a role in energy metabolism, upregulated in numerous cancers<sup>82,83</sup> |
| *rab-6.2* | *RAB6A* | Involved in the transport of proteins from Golgi body to endoplasmic reticulum<sup>84</sup> |
| *sem-5* | *GRB2* | Activates RAS/MAPK downstream of the EGFR receptor<sup>85</sup> |
| *R119.3* | *DHRS2* | Member of the short chain dehydrogenases family, metabolizes steroid hormones and lipids<sup>86</sup> |
| *pak-1* | *PAK1* | Kinase involved in cytoskeletal remodeling, contributes to metastasis of many cancers<sup>87,88</sup> |
| *capg-1* | *NCAPG* | Part of the condensin complex which condenses chromosomes during mitosis and meiosis<sup>89</sup> |

### CFIm25 regulates migration and invasion of KRAS-mutant cancer cells

To determine if our observations in *C. elegans* were conserved, we assessed *CFIm25* function in human KRAS-mutant cancer cells. We employed two HCT116 isogenic cancer cell lines, where the parental line had a heterozygous KRAS G13D mutation (KRAS G13D) and its isogenic counterpart had the mutant KRAS allele knocked out (KRAS WT)^47^. Using these cell lines, we explored the mechanism by which CFIm25 functions in the context of mutant Ras in human cancer cells.

Ablation of CFIm25 by siRNAs resulted in spindle-like morphology and cytoplasmic protrusions in cells **(Supplementary Fig. 5a)**. This was reminiscent of cells undergoing an epithelial to mesenchymal transition (EMT), which is associated with metastatic characteristics such as increased migration and invasion. Indeed, the depletion of CFIm25 significantly increased the migratory and invasive properties of KRAS G13D cells (**Fig. 3a**). Compared to KRAS G13D cells, KRAS WT cells showed reduced migration and invasion, and depletion of CFIm25 in KRAS WT cells did not significantly alter their migratory or invasive potential (**Fig. 3a**). We corroborated the EMT phenotype using markers and observed a decrease in the epithelial marker E-cadherin and an increase in expression of the mesenchymal marker Vimentin in CFIm25-knockdown conditions for both KRAS WT and KRAS G13D cell lines (**Fig. 3b, c**). This suggests that even in the absence of oncogenic mutant KRAS, CFIm25 depletion is capable of causing EMT-like alterations in cells, however co-operation with mutant KRAS is required for increased migration and invasion. Furthermore, we found that the increased migration and invasion of CFIm25-knockdown cells were not associated with increased proliferation. Cell growth and cell cycle analysis revealed that depletion of CFIm25 in KRAS G13D cells led to decreased cell growth and increased fraction of cells in the S and G2/M phases (**Supplementary Fig. 5b, c**), suggesting a shift from proliferation to EMT and increased cell migration.

**Fig. 5.**
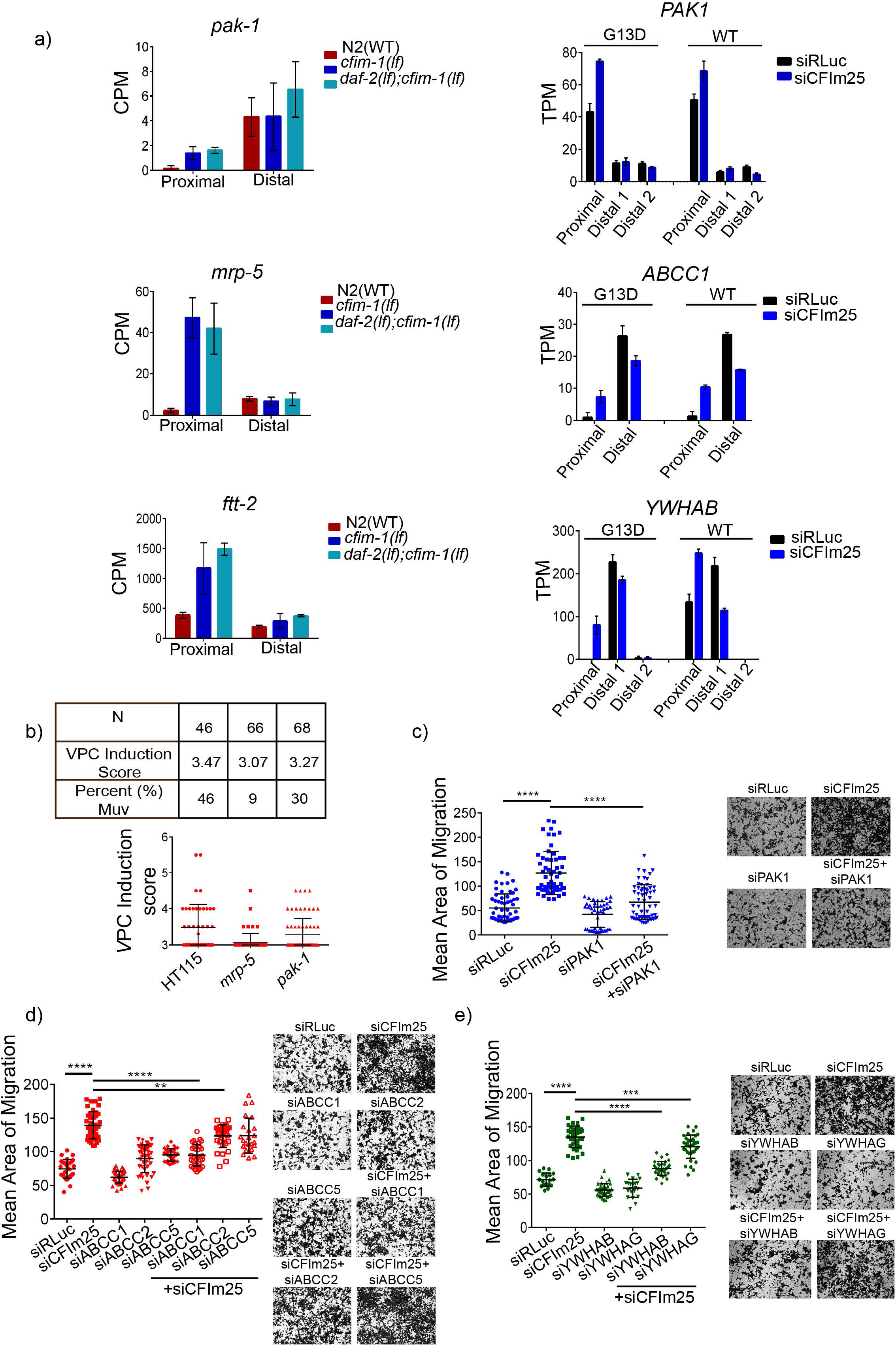
Conserved transcripts regulate oncogenic Ras activity and tumorigenesis. a) Quantification of isoform changes of *pak-1/PAK1*, *mrp-5/ABCC1* and *ftt-2/YWHAB* genes in *cfim-1(lf)*, *daf-2(lf); cfim-1(lf),* and CFIm25-depleted KRAS G13D cells. CPM is Counts per Million and TPM is Transcripts per Million. b) VPC induction scores of *let-60(gf);daf-2(lf);cfim-1(lf)* mutants on RNAi against *mrp-5* and *pak-1*. Percent (%) Muv indicates percentage of animals with more than 3 VPCs induced. N indicates the number of animals measured. Y-axis origin is set to 3 to represent the normal number of VPCs that are induced to divide c) Quantification of migration of KRAS G13D cells with either single depletion or co-depletion of *CFIm25* and *PAK1*. Each data point represents quantification of a field of view acquired across three independent experiments. Representative images are shown for each condition. Validation of knockdown is shown in Supplementary Fig. 8a d) Quantification of migration of KRAS G13D cells with either single depletion or co-depletion of *CFIm25* and *ABBC1/2/5*. Each data point represents quantification of a field of view acquired across two independent experiments. Representative images are shown for each condition. Validation of knockdown is shown in Supplementary Fig. 8a e) Quantification of migration of KRAS G13D cells with either single depletion or co-depletion of *CFIm25* and *YWHAB/G*. Each data point represents quantification of a field of view acquired across two independent experiments. Representative images were shown for each condition. Validation of knockdown is shown in Supplementary Fig. 8a ****p<0.0001, ***p<0.001, **p<0.01, One-way ANOVA with Tukey’s post hoc test

We extended our studies to another human colon cancer cell line DLD-1 with KRAS G13D mutation and three pancreatic cancer lines BXPC3 (KRAS WT), CFPAC-I (KRAS G12V), and HPAF-II (KRAS G12D)^48^. Consistent with our results in HCT116 cells, CFIm25-knockdown increased migration of KRAS-mutant DLD-1, CFPAC-I and HPAF-II lines, but had no detectable effects on the KRAS WT BXPC3 cell line (**Supplementary Fig. 6a, b**). Altogether, our data indicate that CFIm25 depletion has a common role in Ras-driven cancer cells.

**Fig. 6.**
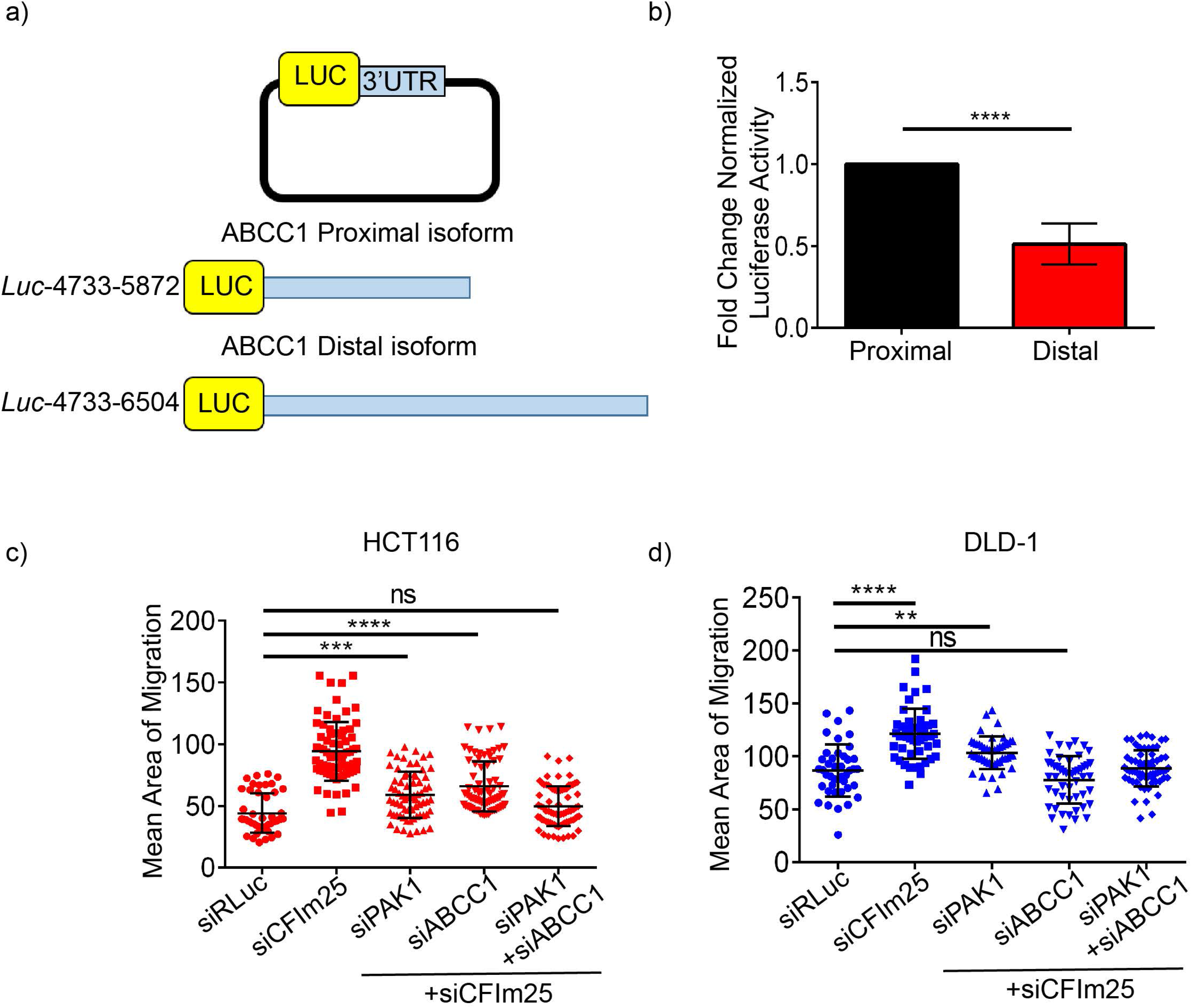
3’UTR sequence elements mediate post-transcriptional regulation of ABCC1. a) Schematic representation of ABCC1 3’UTR luciferase reporter plasmids. Numbers denote the beginning and end of ABCC1 proximal or distal 3’UTR sequences. b) Quantification of luciferase activity for proximal and distal constructs normalized to Renilla luciferase (internal control). c) Quantification of migration of HCT116 KRAS G13D cells with either single, double or triple knockdown for *CFIm25* and/*or PAK1* and *ABCC1*. Each data point represents quantification of a field of view acquired across three independent experiments. d) Quantification of migration of DLD-1 KRAS G13D cells with either single, double or triple knockdown for *CFIm25* and/or *PAK1* and *ABCC1*. Each data point represents quantification of a field of view acquired across two independent experiments ****p<0.0001, ***p<0.001, ns=not significant, One-way ANOVA with Tukey’s post hoc test

### CFIm25 regulates 3’UTR shortening of conserved genes

To identify specific transcripts regulated by CFIm25 in cancer cells we subjected both KRAS mutant and WT HCT116 cell lines with/without CF Im25 knockdown to RNA sequencing (**Fig. 4a**). Sequencing was performed at high depth using Quantification of Alternative Polyadenylation (QAPA) pipeline to quantify the relative abundance of short and long 3’UTR isoforms^49^. We classified the genes into categories based on whether there was an isoform switch (increase in short accompanied by decrease in long or vice versa), changes only in the short (proximal) isoform, or changes only in the long (distal) isoform (**Fig. 4b**). From this analysis, 415 genes were identified with increased expression of shortened isoforms in CFIm25-depleted KRAS mutant cells. We also assessed the APA landscape of KRAS WT cells and found that CFIm25 depletion promoted shortening of 421 genes, of which the 3’UTRs of 165 genes were commonly shortened in both the KRAS mutant and WT isogenic HCT116 cell lines (**Fig. 4c**). We also observed that CFIm25 itself is subject to differential APA regulation in KRAS G13D and WT cells, with WT cells expressing increased levels of the proximal isoform of CFIm25 (**Supplementary Fig. 7a**). This alludes to the possibility that activated KRAS influences expression of CFIm25 in order to modulate APA of specific genes.

**Figure 7.**
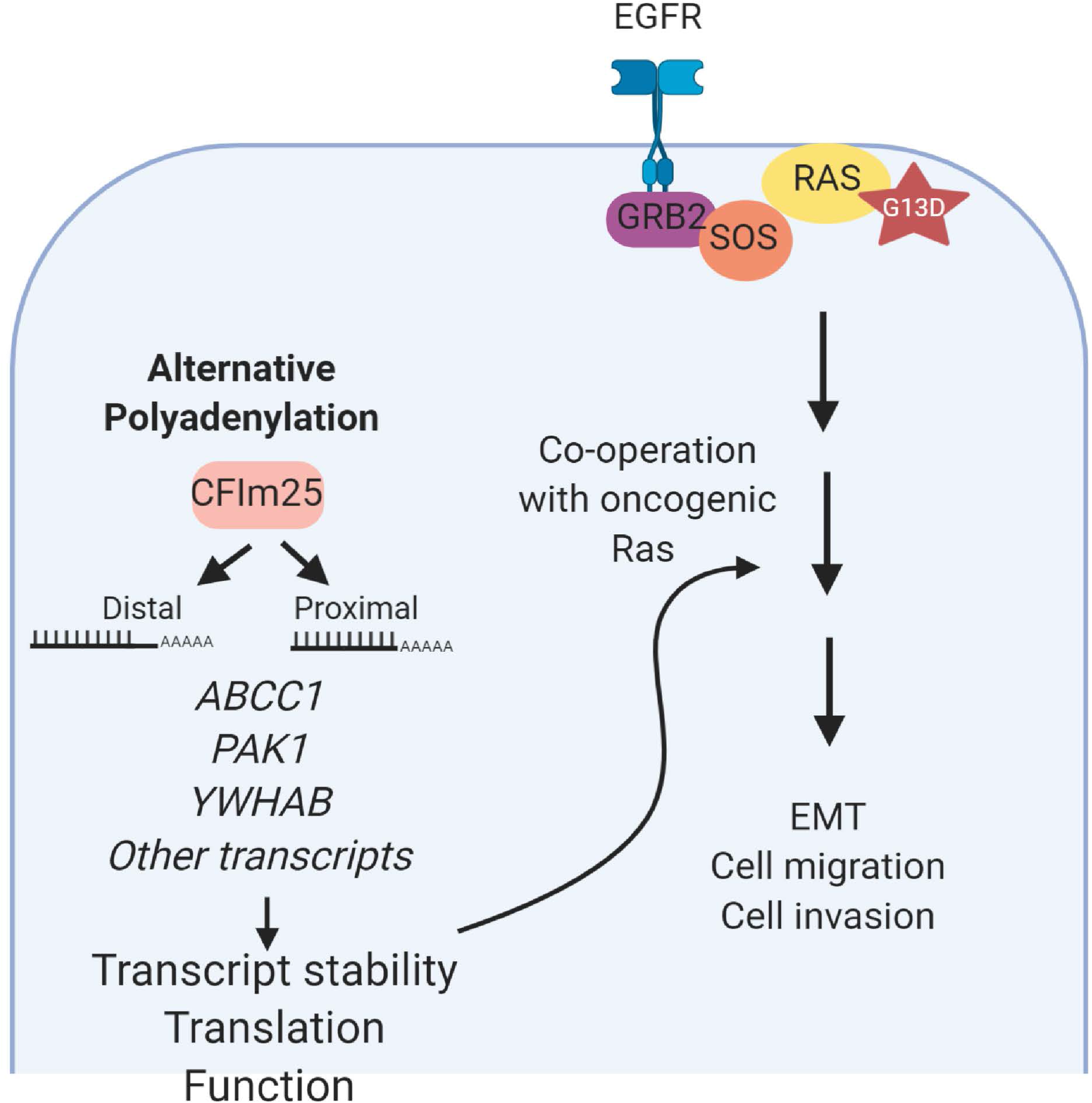
Schematic model of CFIm25 co-operation with oncogenic Ras. CFIm25 regulates APA of multiple genes such as *ABCC1, PAK1 and YWHAB*. Loss of CFIm25 promotes 3’UTR shortening of these genes which in turn can affect the transcript stability, translation and localization and co-operate with oncogenic Ras to regulate EMT, cell migration and invasion

Ingenuity Pathway Analysis (IPA) of shortened transcripts in KRAS G13D cells revealed enrichment in cancer-related signaling pathways, such as protein kinase A, WNT/β-catenin and ERK/MAPK axes, supporting our results that CFIm25 depletion biases APA to shorter isoforms in numerous genes that regulate migration and invasion (**Fig. 4d**). In addition, mining of our RNAseq dataset revealed expression changes in EMT-related genes upon CFIm25 depletion, especially in the KRAS mutant cells, including increased TGF-β2 expression, reduced Occludin, and significant upregulation of multiple EMT-associated chemokines (**Supplementary Fig. 7b**), providing more evidence that CFIm25 regulates EMT^50–52^.

Our screen of genes with shortened 3’UTRs in *C. elegans* uncovered 11 that were required for enhanced Ras signaling in *cfim-1* mutants. We aimed to extend this analysis to conserved genes by comparing the *C. elegans* APA gene list with the human dataset. The 679 *C. elegans* genes shortened in *daf-2(lf); cfim-1(lf*) double mutant were converted to their corresponding human orthologs using Ortholist 2 (**Fig. 4e**)^53^. This generated a list of 878 human genes (a single *C. elegans* gene can have multiple orthologs in humans). We performed IPA analysis on these genes and observed enrichment in similar cancer-related pathways as the human gene set (**Fig. 4e**). Comparing the *C. elegans* list to the human dataset revealed 28 *C. elegans* genes that had 33 orthologous human counterparts (**Fig 4f, Table 2**), including the genes that suppressed Muv such as *pak-1/PAK1*, as well as a number of genes that did not fall into the functional categories we initially focused on. Ablation of the 28 *C. elegans* genes by RNAi in *let-60(gf)*; *daf-2(lf)*; *cfim-1(lf)* triple mutants uncovered genes that either supressed or enhanced the Muv phenotype (**Fig. 4g**). In addition to *pak-1/PAK1,* other genes that supressed the Muv phenotype were *mrp-5/ABCC*, *hst-1/NDST1* and *mml-1/MLXIP*, which have pro-survival or chemo-resistance roles in cancers^54–56^. In summary, these results highlight the conserved role that *cfim-1*/CFIm25 plays in regulating 3’UTR shortening of specific transcripts that modulate Ras signaling.

**Table 2.**
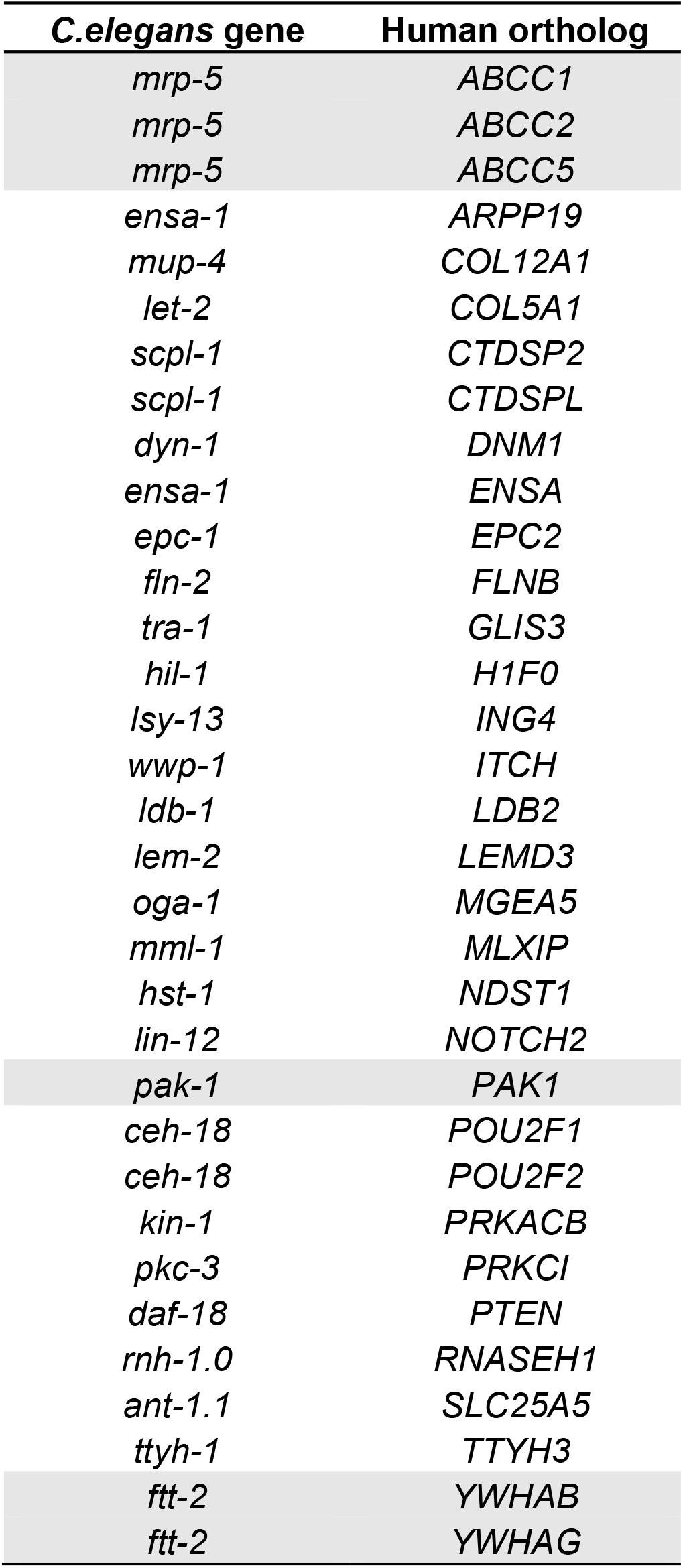
Genes that are commonly regulated by 3’UTR shortening in *C. elegans* and human cancer cells.

### Multiple APA-regulated transcripts contribute to CFIm25-dependent cell migration

To assess the importance of conserved genes in the context of CFIm25-dependent Muv and cell migration, we focused on *pak-1*/PAK1, *ftt-2*/*YWHAB/G*, and *mrp-5*/*ABCC1/2/5* because they have been strongly implicated in cancer biology, and the precise mechanisms for their transcriptional regulation are not known **(Table 2).***cfim-1/CFIm25* depletion increased the proximal isoform expression of these genes in both *C. elegans* and HCT116 cancer cells **(Fig. 5a)**.

To corroborate our observations in Fig. 4g, we ablated *pak-1* and *mrp-5* in *C. elegans let-60(gf); daf-2(lf); cfim-1(lf)* triple mutants and quantified VPC induction. Individual depletion of *pak-1* or *mrp-5* supressed Muv and reduced VPC induction scores from 3.47 to 3.27 and 3.07, respectively (**Fig. 5b**). Ablation of *ftt-2* caused developmental arrest that prevented analysis of vulva development. Co-suppression of *PAK1*, *YWHAB* or *ABCC1* with *CFIm25* significantly repressed migration of HCT115 cells compared with CFIm25 depletion, while knockdown of the APA-regulated genes alone had only subtle effects (**Fig. 5c, d, e, Supplementary Fig. 8a**). Genes within the same family, such as *YWHAG* and *ABCC2/5* supressed the CFIm25-dependent migratory phenotype to a lesser extent (**Fig. 5d, e**). These genes, with the exception of *PAK1*, did not show overall changes in mRNA levels after CFIm25 knockdown (**Supplementary Fig.8b**), highlighting the contribution of specific isoforms to KRAS-dependent cell migration. Although genes such as ABCC1 displayed 3’UTR shortening in CFIm25 depleted KRAS WT cells, these cells did not exhibit increased migratory properties, indicating that CFIm25-dependent APA of transcripts such as ABCC1 cooperate specifically with oncogenic KRAS.

### 3’UTR shortening mediates Post-transcriptional regulation of ABCC1

To better understand how APA of *ABCC1* collaborates with oncogenic Ras, we assessed the impact of the specific 3’UTR isoforms on post-transcriptional regulation of ABCC1. The short and long 3’UTRs were cloned into a luciferase reporter construct (**Fig. 6a**). Luciferase activity was significantly reduced for the ABCC1 distal construct compared to the proximal construct, indicating that the long 3’UTR contains sequence elements that can reduce protein output, either through RNA degradation or translation. Therefore, loss of CFIm25 leads to 3’UTR shortening of *ABCC1* and escape from negative regulation by sequence elements present in the long 3’UTR (**Fig. 6b**).

### PAK1 and ABCC1 cooperate in regulating CFIm25-mediated cell migration

Our analysis identified multiple transcripts required for CFIm25-mediated migratory effects. However, ablation of just one transcript in the context of CFIm25 depletion was not able to completely suppress migration to control levels in HCT116 cells. Therefore, we asked if co-ablation of *PAK1* and *ABCC1* would have a stronger effect than depletion of either gene alone. Triple knockdown of *CFIm25*, *PAK1* and *ABCC1* supressed cell migration to control levels (**Fig 6c**), indicating cooperation of genes subject to APA in oncogenic KRAS output. We also assessed the importance of these transcripts in the DLD-1 cell line and found that ablation of *ABCC1* alone was sufficient to inhibit migratory effects to control level after CFIm25 depletion, but with less dependency on *PAK1* (**Fig. 6d**). This indicates that different cancers and tissues have unique dependencies on distinct sets of APA-modulated genes.

## DISCUSSION

Understanding the cooperation between post-transcriptional regulatory mechanisms and activated oncogenes is an important challenge for identifying cancer dependencies that extend beyond genetic alterations. Here, we show that alternative polyadenylation functions as a conserved process in the modulation of oncogenic Ras output. Specifically, the APA factor *cfim-1/CFIm25* functions to dampen hyperactive Ras-dependent output by biasing 3’UTR usage of multiple transcripts.

Loss of *cfim-1* results in 3’UTR shortening of numerous genes in *C. elegans* that increase the Muv phenotype with oncogenic gain-of-function mutations the Ras orthologue *let-60*. Interestingly, loss of *cfim-1* alone was insufficient to induce the Muv phenotype. Instead, *cfim-1* ablation enhanced Muv when the strong *let-60(n1046)* allele was suppressed by a reduction-of-function allele in the insulin-like growth factor receptor *daf-2*, or in the weaker *let-60(ga89)* single mutant. Similarly, in isogenic HCT116 colon cancer cells, CFIm25 depletion promoted the 3’UTR shortening of key oncogenes and induced EMT-like phenotypes in both Ras mutant and WT cells. However, increased migration and invasion were only observed in cells expressing mutant KRAS, suggesting that these alterations in the APA landscape cooperate with oncogenic Ras to modulate signaling output in both *C. elegans* and human cells. Studies have shown that thresholds of mutant KRAS activity dictate progression of cancer and metastasis^57–59^. We suggest that APA functions to lower threshold for activated oncogene signaling in both organ development and cancer cell migration.

Although CFIm25 depletion has been associated with increased prevalence of cancer phenotypes in different tissues^60–63^, our study provides new insights into how conserved genes such as *pak-1/PAK1*, the *mrp-5/ABCC* family of transporters, and *ftt-2/14-3-3* family that are subject to APA by CFIm25 co-operate with oncogenic Ras. Ablation of these genes supressed the Muv phenotype in worms and migration of human cancer cells caused by *cfim-1*/CFIm25 depletion, indicating that these are conserved determinants of oncogenic Ras output. Of note, a recent study has shown PAK1 undergoes 3’UTR shortening in response to CFIm25 depletion in glioblastomas, and this serves as a marker of poor prognostic outcome in patients^64^. It is possible that the aggressiveness of cancers such as glioblastoma is dependent on cooperation between genes subject to APA with activated oncogenic pathways. Although KRAS is not highly mutated in glioblastoma, the MAPK/ERK pathway is frequently hyperactivated in this disease and is therefore a desirable target for therapeutic intervention^65^.

The role of ABCC1 in cancers has been traditionally associated with multi-drug resistance (MDR) through the efflux of chemotherapeutic compounds such as doxorubicin, etoposide and vincristine^56,66^. High levels of ABCC1 have also been shown to correlate with drug resistance and increased metastasis of breast cancers, neuroblastomas and non-small cell lung cancers^67–70^, although the precise mechanisms of altered ABCC1 expression is unknown. Our study reveals that ABCC1 can also regulate cell migration downstream of CFIm25, supporting oncogenic functions independent of drug efflux. Analysis of the 3’UTR of ABCC1 revealed that the long 3’UTR contains consensus binding sites for microRNA miR-145a, which has been associated with ABCC1 expression in breast cancers as well as sensitivity to chemotherapy^71^. Therefore, in addition to its role in drug resistance, it is possible that 3’UTR shortening of ABCC1 facilitates escape from miR-145 mediated repression and contributes to the increased migratory property of cancer cells.

Since CFIm25 depletion affected alternative polyadenylation and shortening of over 400 transcripts, we hypothesized that multiple transcripts might affect cell behaviors such as migration in a combinatorial manner. Indeed, co-depletion of two APA-regulated transcripts (*ABCC1* and *PAK1*) restored KRAS mutant HCT116 cells to control levels of migration. These observations suggest that APA serves to modulate expression thresholds of genes that function in distinct processes that support Ras-driven cellular behaviours. This provides unique opportunities for exploiting the APA machinery, or drugging specific proteins/pathways affected by APA, to treat various cancers.

Understanding how changes in the APA landscape affect cell behaviour is critical since conventional gene expression analysis methods do not always reflect changes in specific isoforms, particularly differences in the 3’UTRs. A recent study where 3’-end sequencing of patient derived chronic lymphocytic leukemia samples was conducted identified novel tumor suppressor genes altered through APA. These genes are not mutated frequently in cancer but possess strong tumor suppressor functions, highlighting the need to extend cancer-related studies beyond genomic DNA sequencing and standard gene expression analyses^72^. Organisms like *C. elegans* offer genetically tractable complimentary *in vivo* systems to assess the cooperation between activated oncogenes and APA factors as well as rapid screening methods for uncovering relevant genes that enforce oncogenic output in human cancer. Combining the powerful tools of *C. elegans* with cultured human cells could accelerate discovery of therapeutic targets that exploit vulnerabilities in cancer.

## MATERIALS AND METHODS

### Crosses to generate double and triple mutants

All crosses were done by plating 3-5 hermaphrodites with 10-15 males on petri dishes filled with NGM containing a bacterial lawn. Approximately 5 F1 hermaphrodites at the L4 stage of development were then plated onto plates. F2 progeny were collected at the L4 stage of development and singled out onto separate plates. F2 worms were incubated until they had laid eggs, then collected and lysed. The worm lysate was used for PCR reactions to confirm the presence of the desired genetic lesion.

### Scoring of multivulva phenotype

The multivulva phenotype was scored based on the degree of vulva induction. Vulva induction refers to the number of vulval precursor cells (VPCs) induced to form part of the vulva at the L2 stage of development. This is measured by visualizing the VPC daughter cells at the L4 stage of development. Visualization was done using a Leica DMRA2 compound light microscope at 63x magnification. Worms were mounted on glass slides on 4% agarose pads and anesthetized with 20mM Tetramisole. Primary vulval precursor cells have undergone 3 rounds of cell division and comprise 8 daughter cells. Secondary daughter cells have undergone 8 rounds of cell division and comprise 7 daughter cells. Tertiary vulval precursor cells have undergone one round of cell division and comprise two daughter cells. These daughter cells and the pattern they form are easily visualized and measured. A normal vulva results from one primary and two secondary vulva cells. These are given an induction score of 3.0 since three VPCs have been induced. Often worms of the Muv phenotype have tertiary VPCs induced to partial primary and secondary fates. These cells have only 3-4 vulva cell daughters forming a pseudovulva. These are given an induction score of 0.5 as “half” of a vulval cell has been induced in these cases. In addition to induction scores, percentage of Muv worms (%Muv) of various genotypes was calculated. In cases where induction scores were calculated, % Muv was calculated by dividing the number of worms with induction scores over 3.0 by the total number of worms measured. In cases where no induction scores were calculated, % Muv was calculated by scoring all worms with at least one pseudovulva and dividing this number by the total number of adult worms. All worms scored for vulva induction were raised at 20°C with the exception of the *let-60(gf);daf-2(lf);cfim-1(lf)* strain which was raised at 16 °C.

### RNAi feeding protocol

RNAi cultures were obtained from the Source Bioscience *C. elegans* RNAi library. Bacteria were struck onto LB plates containing ampicillin and tetracycline (working concentrations 50mg/mL and 5mg/mL respectively). Plates were incubated overnight at 37°C. Single colonies were picked and inoculated in liquid LB^amp+tet^ overnight at 37°C with shaking. Following the overnight inoculation, isopropyl β-D-1-thiogalactopyranoside (IPTG) was added to each culture (working concentration 0.5μM) and incubated at 37°C with shaking for 4 hours. The bacterial culture was then concentrated 2x or 10x by centrifugation and mixing. The culture was then plated on solid NGM media with added carbenicillin and IPTG (working concentrations 25μg/mL and 25mM respectively). Worms were added to these RNAi plates at the L4 stage and measurements were taken of the F1 progeny.

### Candidate RNAi screening protocol

For the screening of DAF-16 targets, single colonies of each RNAi from the Source Bioscience C. elegans RNAi library were inoculated into liquid LB media and induced with IPTG as described in the “RNAi feeding protocol” section. Bacterial cultures were plated onto 12 well plastic plates filled with solid NGM RNAi media (as described in “RNAi feeding protocol”) so that each well contained a different RNAi bacteria strain. Each plate also included one well with negative control bacteria HT115, and positive control bacteria expressing *daf-16* RNAi. Bacterial lawns were allowed to grow overnight at 37°C. The next day two L4 *daf-2(e1370); let-60(n1046)* worms were picked into each well. Worms were allowed to grow until pseudovulvae could be seen with confidence in the *daf-16* RNAi positive control well. Plates were then scanned through the rest of the wells, noting RNAi treatments that resulted in worms being noticeably more Muv than the negative controls, and appearing similar to worms treated with *daf-16* RNAi. Each plate was replicated and a positive hit was only scored if a Muv phenotype could be seen in both replicates. Positive hits were subjected to a more stringent analysis of ability to induce Muv. We grew up more cultures of promising RNAi strains and plated them 2x concentrated onto 10cm solid NGM RNAi plates. We plated 3-4 *daf-2(e1370); let-60(n1046)* L4 worms onto these plates and scored 15-20 F1 progeny for vulva induction under 63x magnification.

For the screening of 3’UTR shortened genes, RNAi cultures were grown overnight and induced with IPTG for 4 hours. Cultures were then concentrated 10X and plated onto NGM plates. 4-*5 let-60n1046(gf), daf-2(lf); cfim-1(lf)* L4 worms were picked onto each RNAi plate and percent Muv was calculated for the F1 progeny.

Only those RNAi treatments that were able to produce a significant increase in Muv induction compared to the negative controls HT115 were counted as hits.

### Generation of CRISPR strains

The *cfim-1(lf)* CRIPSR strain were created with CRISPR technology using the self-excising drug selection cassette as previously described^90^. The “aaacctccg” sequence at position 79 from the *cfim-1* start codon was mutated by deleting 5 bp “aaacc” and changing “tccg” to “tagt” to create frame shift and a premature stop codon (TAG).

### RNA extraction

For *C.elegans* samples, RNA was extracted from L4 stage worms using TRIzol (Invitrogen). RNA from human cancer cells was extracted 48h post transfection using the RNeasy Kit from Qiagen according to the manufacturer’s instructions.

### RNA-seq library preparation and sequencing

To construct RNA-seq libraries from *C.elegans* samples, we used an automated QuantSeq 3’mRNA-seq (Lexogen GmbH, Vienna) using the Agilent NGS Workstation (Agilent Technologies, Santa Clara) as per manufacturer’s protocol. Briefly, 250 ng of total RNA was used to generate cDNA. cDNA was amplified with 17 PCR cycles as determined by qPCR analysis using the PCR Add-on kit (Lexogen). ERCC RNA Spike-ins were added following manufacturer’s instructions. The resulting libraries were quantified with Qubit DNA HS (Thermo Fisher, Waltham) and fragment sizes analyzed on the Agilent Bioanalyzer using the High Sensitivity DNA assay prior to sequencing. Sequencing was performed at The Centre for Applied Genomics (Toronto, Canada) across 2 lanes of an Illumina HiSeq2500 rapid run flowcell (Illumina, San Diego) with SR100 bp.

For sequencing of human cancer cell samples, total RNA was treated with the DNA-free DNA Removal Kit (cat # AM1906; Thermo Fisher Scientific Inc., Waltham, USA) to remove contaminant DNA. DNase-treated total RNA was then quantified using Qubit RNA BR (cat # Q10211, Thermo Fisher Scientific Inc., Waltham, USA) fluorescent chemistry and 1 ng was used to obtain RNA Integrity Number (RIN) using the Bioanalyzer RNA 6000 Pico kit (cat # 5067-1513, Agilent Technologies Inc., Santa Clara, USA). Lowest RIN was 9.1; median RIN score was 9.8. 1000 ng per sample was then processed using the TruSeq Stranded mRNA Library Prep (cat # 20020595; Illumina Inc., San Diego, USA; protocol v. 1000000040498 v00) including PolyA selection, with 8 minutes of fragmentation at 94 °C and 15 cycles of amplification. 1uL top stock of each purified final library was run on an Agilent Bioanalyzer dsDNA High Sensitivity chip (cat # 5067-4626, Agilent Technologies Inc., Santa Clara, USA). The libraries were quantified using the NEBNext Library Quant Kit for Illumina (cat # E7630L, New England Biolabs, Ipswich, USA) and were pooled at equimolar ratios after size-adjustment. The final pool was run on an Agilent Bioanalyzer dsDNA High Sensitivity chip and quantified using the NEBNext Library Quant Kit. The quantified pool was hybridized at a final concentration of 2.2 pM and sequenced on the Illumina NextSeq 500 platform using two High-Output v2 flowcells, for an average of 85 million 1×85 bp reads.

### Analysis of 3’UTRs

#### C.elegans

##### Sequencing read processing

144 base pair single ends sequencing reads were obtained. Reads were trimmed using Cutadapt as follows: 1) first 12bp of reads were trimmed; 2) basepairs with sequencing quality lower than 30 were trimmed from the ends of reads; 3) polyA stretches at the end of reads were trimmed using 10 As as an “adaptor” sequence. Only reads with at least 5 As trimmed at the ends are used for subsequent analysis. ERCC transcripts were only used to ensure library quality and were not included in the subsequent analyses.

##### Polyadenylation (PA) site identification

The 3’ most nucleotide of each read was used to create a track of reads pile-up. R package “derfinder”^91^ (version: 1.12.0) was used to identify expressed regions separately for reads that mapped to the positive or the negative strand. Essentially, any consecutive positions with more than 1 RPM (reads per million mapped reads) were identified as an expressed region. Regions were extended 5bp upstream. Regions that are within 5bp from each other were then merged.

PolyA clusters were then mapped to *C. elegans* gene annotation (WBcel235, obtained from Ensembl 85). Meanwhile, sequence composition downstream of each ER was examined to identify ERs that are potentially originated from internal polyA priming events. Specifically, two criteria are used: 1) within 30bp downstream and 5bp upstream from each ER end, if the pattern of 18As (allowing 5 mismatches) can be found; 2) within 10bp downstream and 5bp upstream from each ER end, if the frequency of As is larger or equals to 60%. In addition, 35bp upstream from each ER end is searched for the most common two polyA signal, AATAAA and ATTAAA.

##### PA site annotation

ERs that map to exons are annotated to the associated genes. ERs that do not map to exons are annotated with the following steps:

1. ERs that are within 3kb from the nearest transcript end site (TES) upstream are annotated to the corresponding gene
2. ERs that map to introns are annotated to the associated genes
3. In the rare case (n=216) that an ER maps both to an intron and downstream of a gene, if the ER maps within 200bp of a gene’s TES, it is annotated to that gene, otherwise it is annotated to the intron-containing gene.

Finally, all annotations are merged resulting in more than 99% of the ERs being annotated to a gene. Number of reads mapped to each ER is then calculated.

##### Identify alternative PA sites per gene

We systematically processed PA sites that mapped to each gene and used stringent criteria to analyze only confident PA sites. We consider a few different situations:

1. PA sites that are likely derived from internal polyAs are discarded even if the PA site is mapped to exons.
2. An exception to 1): If an internal polyA derived site (determined in “PA site annotation”). is the longest potential PA site or the second longest PA site followed by a real PA site, and also contains a significant proportion (more than 50%) of all reads mapped to this gene, it is considered a “valid PA site”. In these two cases, although we cannot precisely pinpoint PA sites (due to internal polyA priming within the 3’ UTR) we have high confidence that this polyA signal is derived from the longest 3’UTR variant (because no signal is detected downstream) and can be used to assay alternative PAs (APAs).
3. If more than 75% of the reads mapped to this gene map to internal polyA-derived sites, this gene is not considered for APA analysis.
4. If potential PA sites are within 15bp of each other, merge them.
5. After merging, only keep the PA sites containing more than 5% of all mapped reads of this gene and have more than 10 reads in at least 3 samples for further analysis. The purpose is to filter out extremely weak PA signals.
6. Finally, genes with more than 1 PA site after the filtering are subjected to DEXSeq analysis.

In total, we have 5793 genes with more than 1 ER mapped. 1236 of them have only 1 PA site after filtering for internal polyA-derived ERs (steps 1 & 2). 91 of them have more than 75% of the reads mapping to polyA-derived ERs (step 3). 2048 genes have only 1 PA site after merging and filtering for low count PA sites (steps 4 & 5). Ultimately, after our stringent filtering, 2418 genes were analyzed for altePA events. For convenience, we order and name the PA sites based on their relative positions along the gene body and “E001” refers to the longest PA site.

##### APA analysis

To characterize genes that undergo APA, we used R package “DEXseq” (version: 1.24.2)^92^ and examined every pair of comparisons. We applied a cut-off of adjusted p value < 0.05 and fold change > 1.5. To summarize the results, we only considered the comparisons between N2 (the wildtype) and cfim1 mutants. For each gene in each comparison, we annotate based on its APA condition. The top 4 primary situations are: 1) the longest PA site is expressed higher in WT; 2) at least one of the shorter PA sites is expressed higher in mutants; 3) the longest PA site is expressed higher in WT and at least one shorter PA site is expressed higher in mutants; 4) the longest PA site is expressed higher in mutants and at least one shorter PA site is expressed higher in the WT. Other more complicated APA cases were not specifically classified and are represented as “Others”.

##### Differential gene expression (DGE) analysis

First, the number of reads mapped to each gene is counted by adding the reads in ERs that mapped to 1) any exons; 2) being annotated to this gene and pass internal polyA filtering criteria (described in “Polyadenylation (PA) site identification”). Next, DGE analysis is performed using R package “DESeq2” (version 1.18.1)^93^. Specifically, count data from all samples were normalized together and pairwise comparisons of conditions of interest were performed with the “ifcShrink()” function. A cut-off of adjusted p value < 0.05 and fold change > 2 was used to identify significantly differentially expressed genes.

##### Human cancer samples

Quantification of 3ʹUTR isoforms: To estimate 3ʹUTR isoform abundances we used the alternative polyadenylation analysis tool QAPA version 1.2.0^94^.In short, QAPA constructs a refined set of 3ʹUTR isoforms from the GENCODE basic gene annotations using external databases of polyadenylation sites^95^. Next, QAPA selects 3ʹUTRs from the most downstream exon of the gene to resolve ambiguities in quantification of 3ʹUTR isoforms with reads generated by RNA-seq. Using this refined 3ʹUTR annotation library, we quantified 3ʹUTR isoforms using raw RNA-seq reads with the alignment-free isoform quantification tool Sailfish version 0.10.0^96^. Based on mRNA quantification from biological replicates per each cell type, we calculated average, standard deviation and coefficient of variation (CV) for each 3ʹUTR isoform.

### Gene Ontology analysis

*C.elegans* genes were classified into biological categories by selecting the GO biological function on Genemania. For overlapping with human dataset, *C.elegans* genes were converted into their human orthologs in Ortholist 2.0^53^. Resulting orthologs were used for subsequent IPA analysis or overlaid with human data set to identify commonly regulated genes. Ingenuity pathway analysis (IPA) was performed on both human as well as worm-converted genes to classify genes into signaling pathways.

### Cell culture

HCT116 KRAS G13D, KRAS WT cells and DLD-1 cells were maintained in McCoy’s 5A medium supplemented with 10% FBS. Pancreatic BX PC3, CFPAC I and HPAF II cell lines were grown in DMEM supplemented with 10% FBS. Cell lines were validated by STR profiling.

### siRNA transfection

For siRNA knockdown experiments, 300 000 cells/well were plated on 6-well plates and transfected with pooled siRNA at a final concentration of 25nM using Lipofectamine RNAiMax according to the manufacturer’s instructions. All siRNAs were obtained from Dharmacon.

### Growth assay

For growth assays, cells were harvested 24h post transfection, counted and 3000 cells/well were plated on 96-well plates. Growth was measured using Presto Blue (Invitrogen) and measuring fluorescence using a plate reader according to the manufacturer’s instructions. Readings were taken at 0, 24, 48 and 72h post-plating for time course experiments.

### Cell cycle analysis

Cell cycle analysis was performed using the Click-iT™ Plus EdU Alexa Fluor™ 647 Flow cytometry assay kit (Thermofisher Scientific). To obtain the cell cycle profile, HCT116 cells +/− CFIm25 knockdown (72h) were pulse labelled with 10μM Edu for 2 hours. The cells were then harvested and fixed by adding ice cold ethanol dropwise to the cells. The cells were then permeabilized, washed and Click-iT™ EdU was detected using the manufacturer’s protocol. Cells were finally suspended in 0.5 mL of FxCycle™ PI/RNase Staining Solution (Invitrogen). Samples were analysed on a Fortessa flow cytometer.

### Cell migration and Invasion assays

Cells were harvested 24 post transfection, counted and 80000 cells were plated on top of a Transwell 8μM insert in low (0.1% FBS) media. Complete media (10%FBS) was placed in the well below the insert. Each condition was performed in triplicate. Cells were allowed to migrate for 20 hours before fixing and staining with 1% Crystal Violet in 20% Methanol for 20 mins at room temperature. The top layer of the transwell membrane was scraped to remove excess stain and washed with PBS. Inserts were allowed to dry before imaging on a Nikon Epifluorescence microscope using a 10X objective. Atleast 5 fields of view were acquired per insert. Area of migrated cells was calculated on ImageJ and represented as Mean Area of migration or Mean Area of Invasion. For invasion assays, 150μg/cm^2^ of Matrigel was added to the top of each insert and allowed to solidify before plating cells. Fixation and imaging was performed as described above.

### Immunoflourescence

Cells were transfected for 24h and then harvested and 100000 cells were plated on poly-L-lysine coated glass coverslips for a further 48h. Cells were fixed in 4% paraformaldehyde, permeabilized with 0.1% Triton-X/PBS for 5 mins and blocked for 1 hour with 10% Goat serum/1 % Bovine serum alumin solution. Coverslips were then incubated overnight at 4 degrees with primary antibody-anti-Vimentin (Biolegend). Coverslips were washed and stained with secondary antibody, Goat anti-chicken Alexa 488, Alexa 568 Phalloidin and DAPI for 1 hour at room temperature. Coverslips were washed and mounted using ProLong Gold Antifade Mountant. Cells were imaged on a Leica Lightsheet confocal microscope using the 63X objective.

### Western Blotting

Cells were lysed in 1% NP-40 lysis buffer containing 5 M NaCl, 10% NP-40, 1 M Tris (pH 8.0) supplemented with protease and phosphatase inhibitors. Lysates were cleared by centrifuging at maximum speed. Protein concentration was determined by using the BCA assay (Thermo Fisher Scientific). 30μg of protein was loaded and run on <u>10% Mini-PROTEAN® TGX™ Precast Protein Gels</u> (Bio-Rad). Separated proteins were transferred onto Nitrocellulose membranes (Bio-Rad) and blocked with 5% BSA for 1 hour, followed by incubation with primary antibodies overnight at 4°C. Following three washed with 0.1% Tween-20 in TBS (TBST), membranes were incubated with HRP-conjugated secondary antibodies for 1 hour at room temperature. Blots were washed and developed using ECL substrate (Thermo Fisher Scientific). The following antibodies were used: anti-NUDT21, 1:1000 (Proteintech), anti-Tubulin 1:5000 (Sigma), anti-E-Cadherin 1:1000 (Cell Signaling Technologies), anti-PAK1 1:1000 (Cell Signaling Technologies), anti-Vinculin 1:5000 (EMD Millipore)

### qRT-PCR

cDNA was reverse transcribed using the SuperScript III Reverse Transcriptase kit (Thermo Fisher Scientific) and quantitative PCR was performed using the ssoAdvanced Universal SYBR Green Supermix (Bio-Rad). Primer sequences are included in the table below.

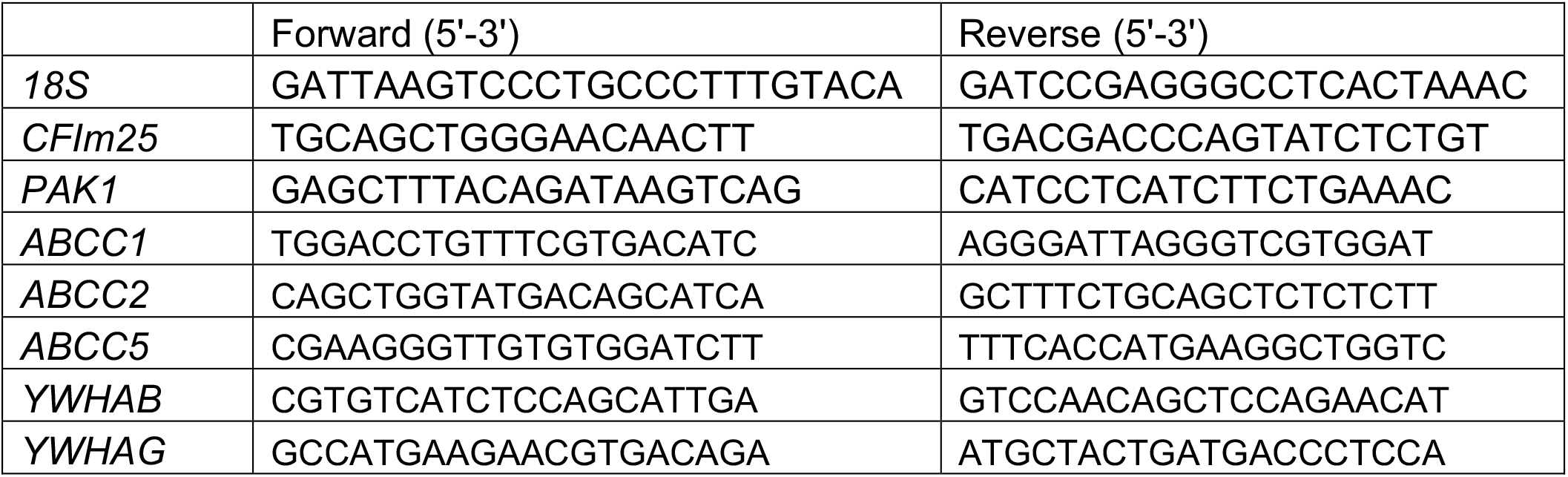

### Luciferase assays

The long and short 3’UTRs cloned into the dual-luciferase vector miTarget vector was obtained from GeneCopoeia. HCT116 cells were plated on 6-well plates and transfected with 50ng of each plasmid. Luciferase expression was measured after 48 hours using the Dual Luciferase Assay System (Promega) according to the manufacturer’s protocol.

## Data Availability

*C. elegans* and mammalian cell line RNA sequencing data generated in this study have been uploaded onto the ArrayExpress database under the accession numbers: E-MTAB-9147 and E-MTAB-9207. Scripts used in data analysis are available upon request.

## Statistical Analysis

All statistical analysis was performed as indicated on Graphpad Prism.

## Creation of Schematic representations

All schematic representations were created with BioRender.com

## Acknowledgements

We thank Dr David Kaplan and Dr. Sean Egan for critical reading of the manuscript and suggestions. We thank the Donnelly Sequencing Centre at the University of Toronto for performing the RNA sequencing. MW is supported by the Canada Research Chairs Program and an Early Researcher Award from the Ontario Ministry of Research and Innovation; KY was supported by a Sickkids Restracomp Fellowship; and HH was supported by CIHR grant (Derry/Daugard). The automated 3’UTR-seq technology development was supported by Genome Canada Disruptive Innovations in Technology Grant to MW. J.E. is funded by Canadian Institutes for Health Research (PJT-148746). W.B.D. and M.D. were supported by the Canadian Institutes for Health Research (PJT 155928). W.B.D. is a Canada Research Chair in Genetic Models of Human Disease and is Vice Chair of Fundamental Research in the Garron Family Cancer Centre at the Hospital for Sick Children.

## SUPPLEMENTARY FIGURE LEGENDS

**Fig. 1.**
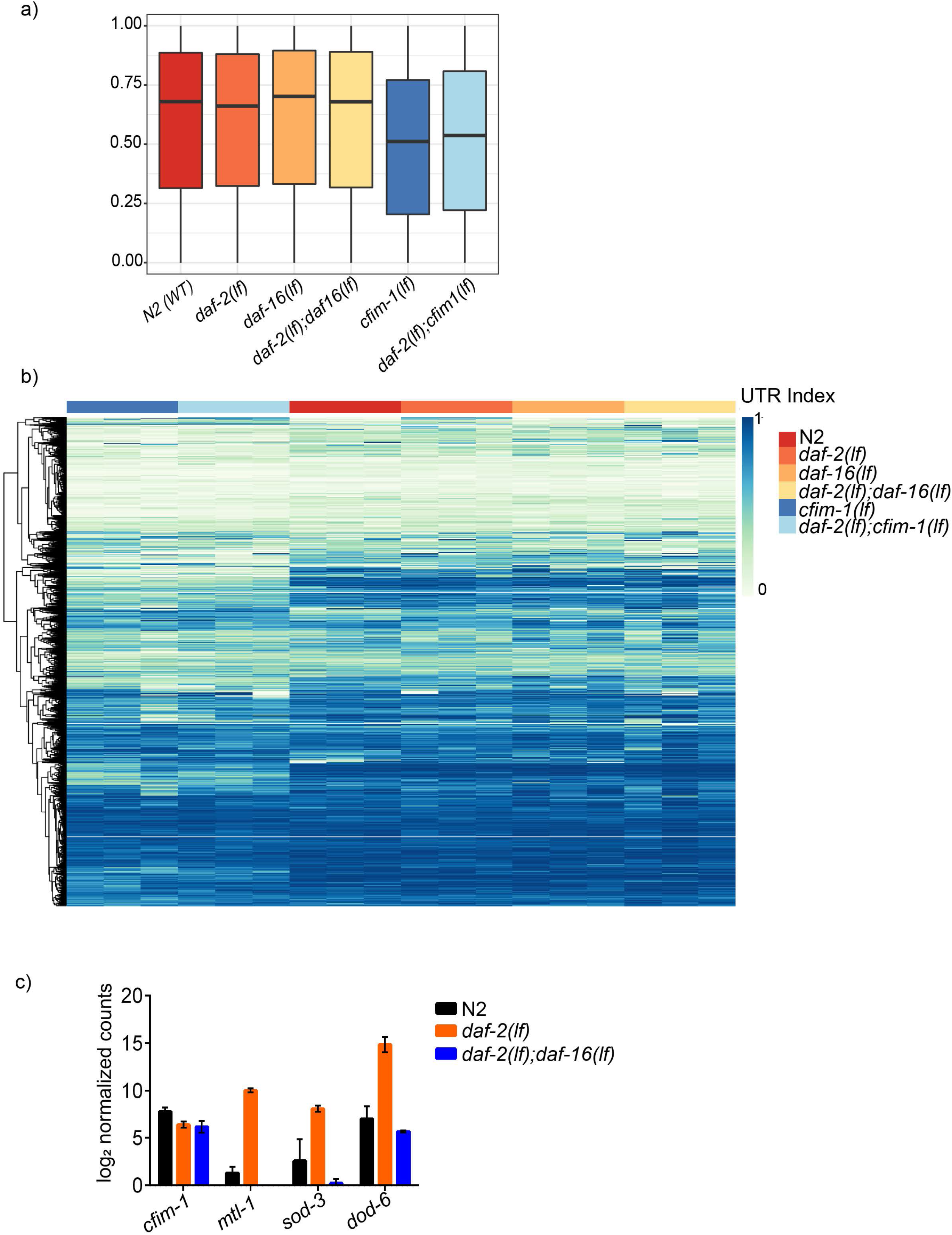
a) Quantification of 3’UTR index (proportion of reads mapping to the longest polyadenylation site) for each gene for N2(WT) and all mutant strains b) Heatmap of UTR indexes for each detected gene across all mutant strains c) Comparison of *cfim-1* expression (log_2_Normalized Counts) in N2, *daf-2(lf)* and *daf-2(lf);daf-16(lf)* mutants. No significant changes in *cfim-1* expression was observed as compared to established DAF-16 targets:*mtl-1,sod-3* and *dod-6*

**Fig. 2.**
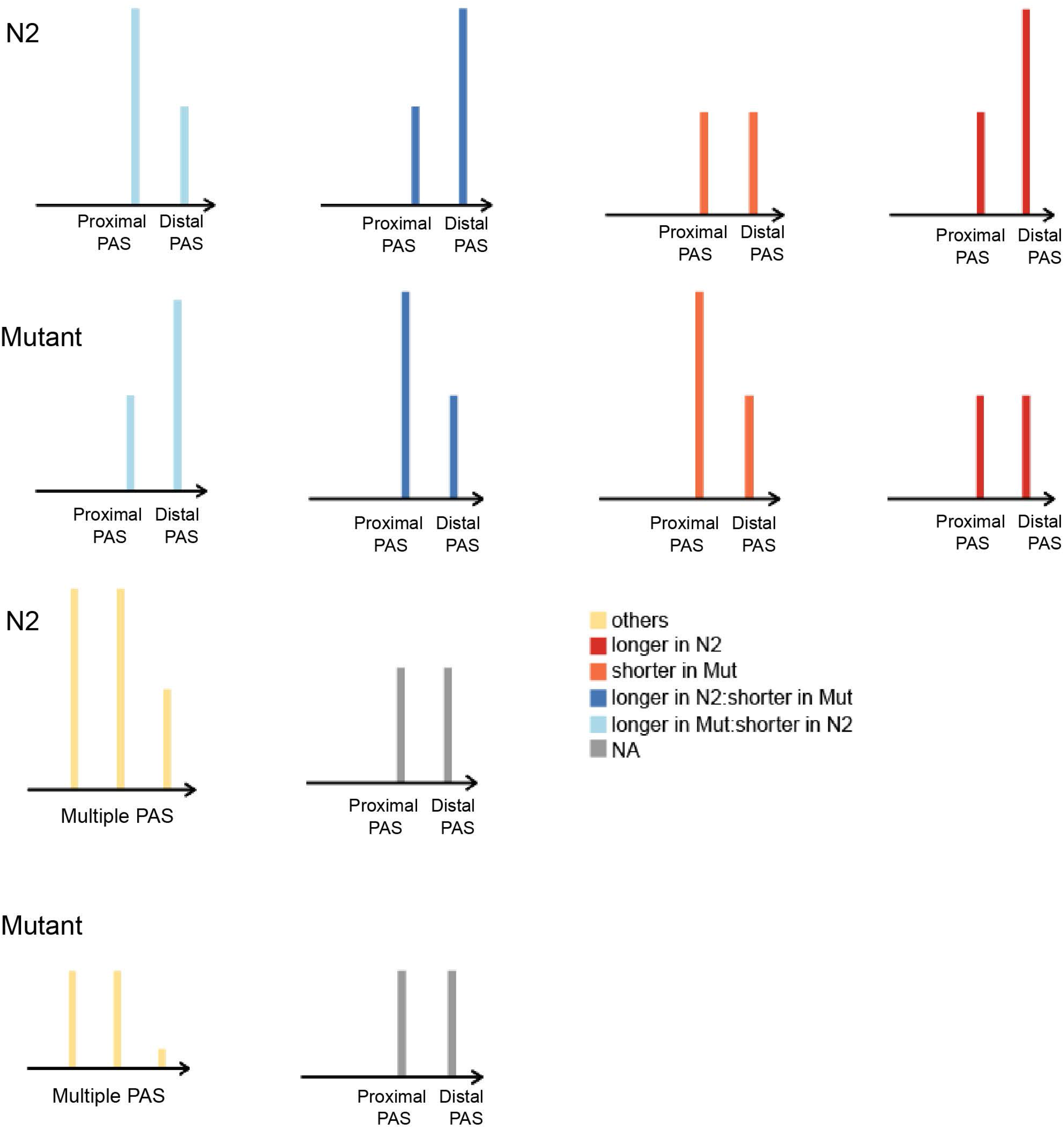
Schematic description of classification of APA events following DEXSeq analysis.

**Fig. 3.**
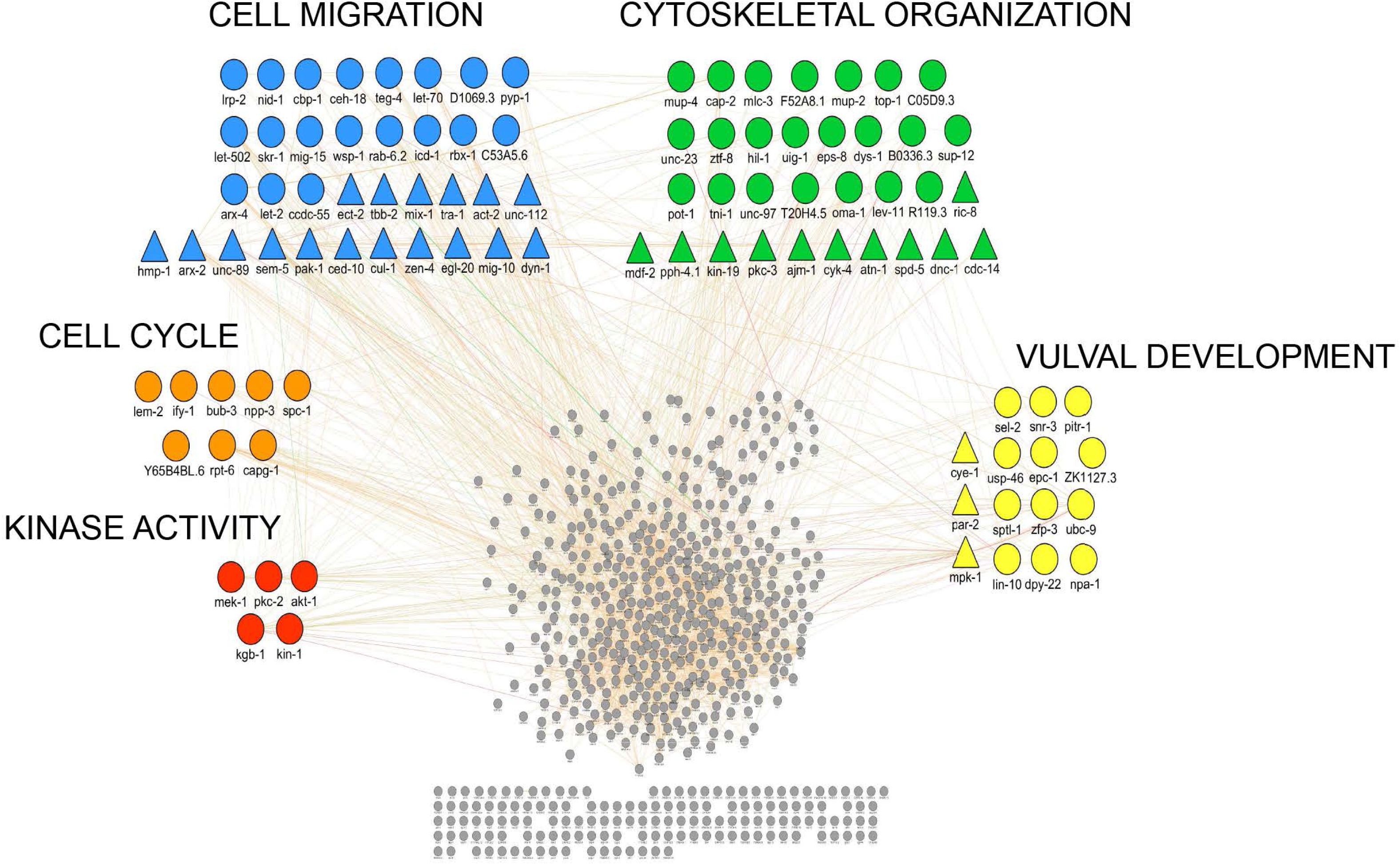
Network analysis of genes which displayed increased expression of short isoforms in *cfim-1(lf)* and *daf-2(lf);cfim-1(lf)* was performed using GeneMania app in Cytosape with gene clusters showing enrichments in Ras related signaling pathways or processes.

**Fig. 4.**
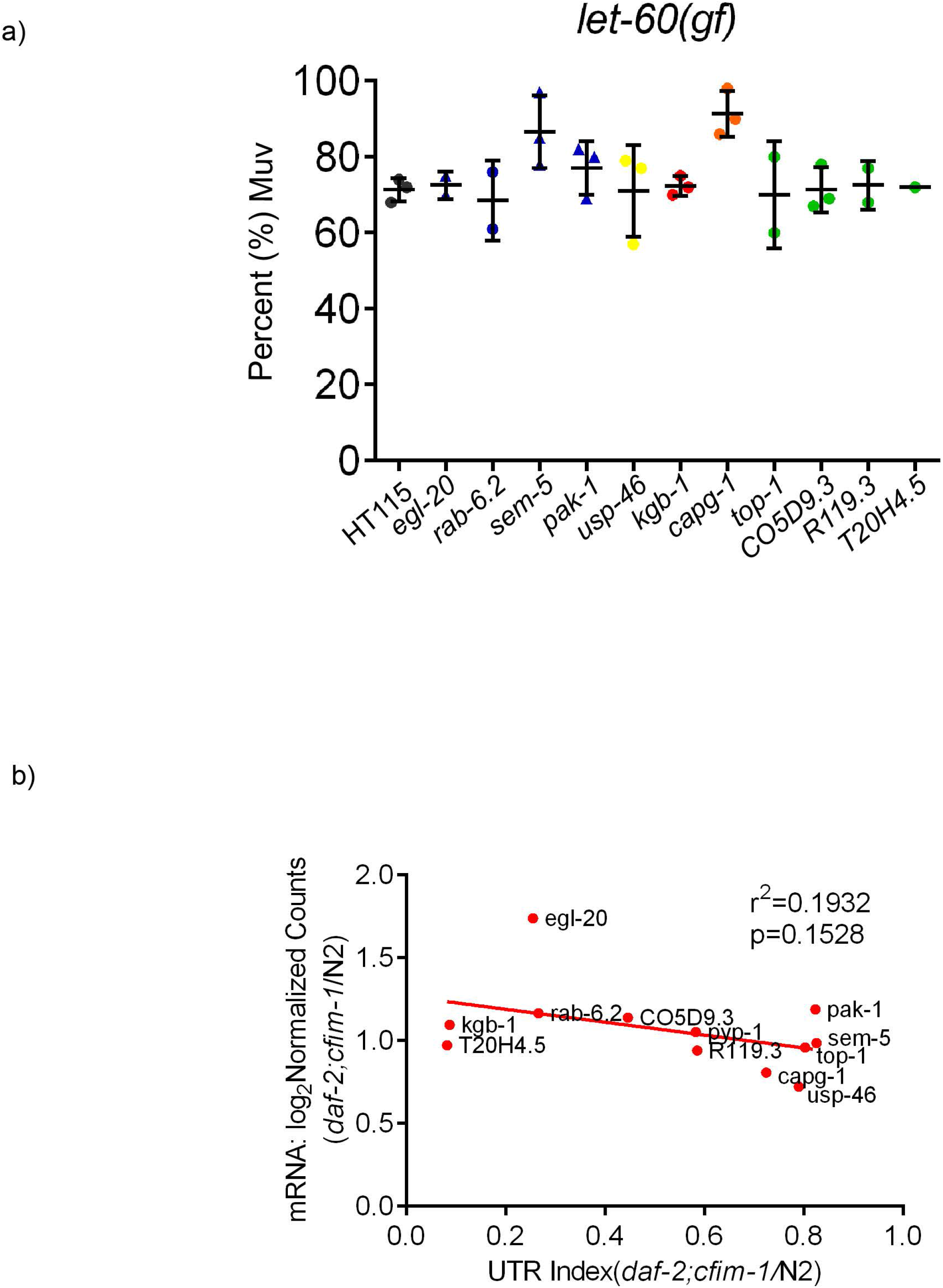
(a) Quantification of percent MUV of *let-60n10469(gf*) mutants on RNAi against shortened genes Each data point represents an independent experiment. Colours and shapes of data points correlate with shapes/colours of genes in network analysis (Supplementary Fig. 3) (b) A correlation analysis (p=0.15) comparing UTR index to total mRNA levels (log_2_ Normalized Counts) for each gene

**Fig. 5.**
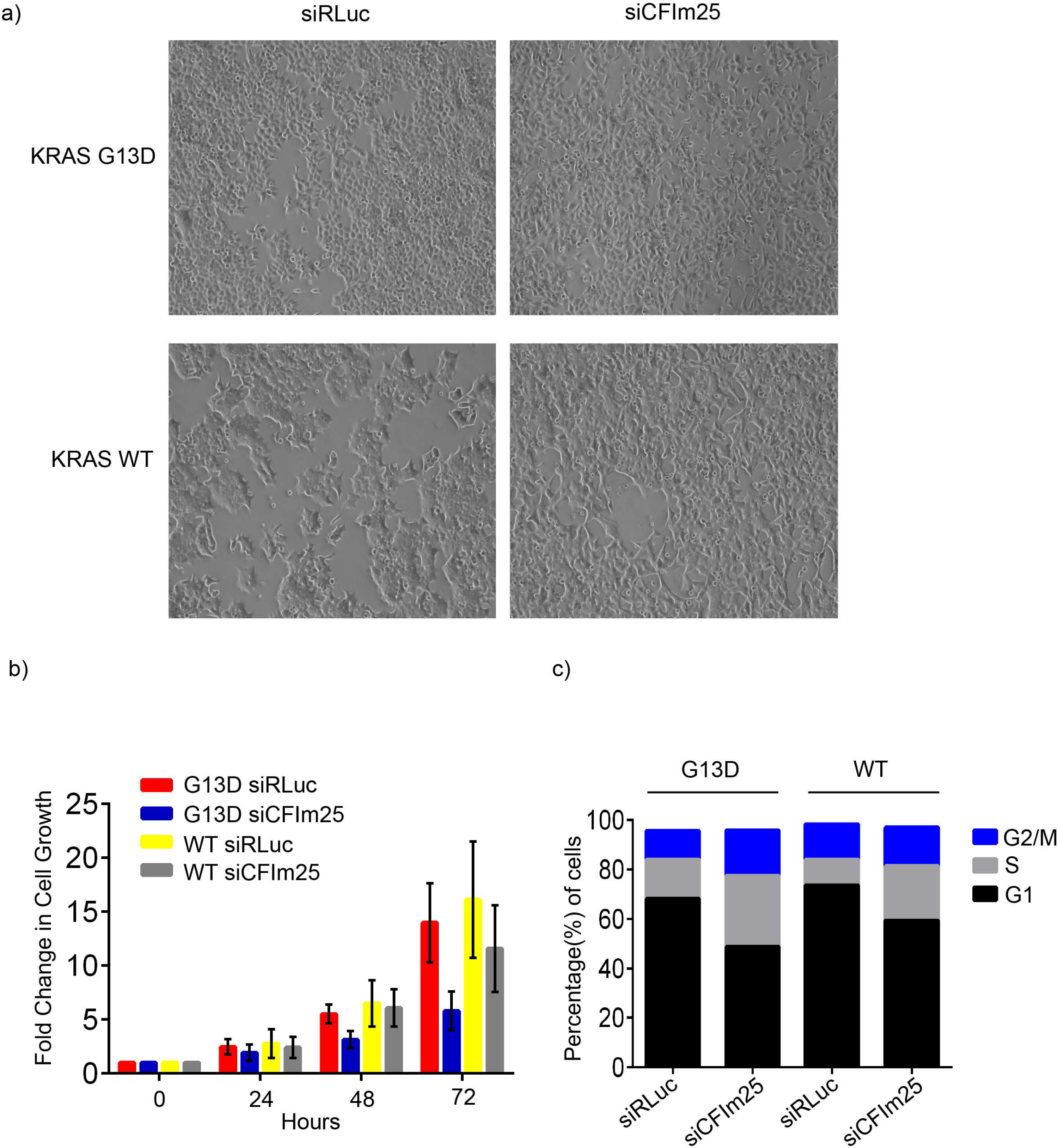
a) Representative bright-field images of isogenic HCT116 cell lines with/without CFIm25 knockdown. b) Time course assay measuring growth of KRAS G13D and KRAS WT with or without CFIm25 knockdown. n=6 independent experiments, mean ± SD. c) Cell cycle analysis of KRAS G13D and KRAS WT with or without CFIm25 after 72h knockdown showing that CFIm25 depletion promotes the accumulation of cells in S and G2 phases.

**Fig. 6.**
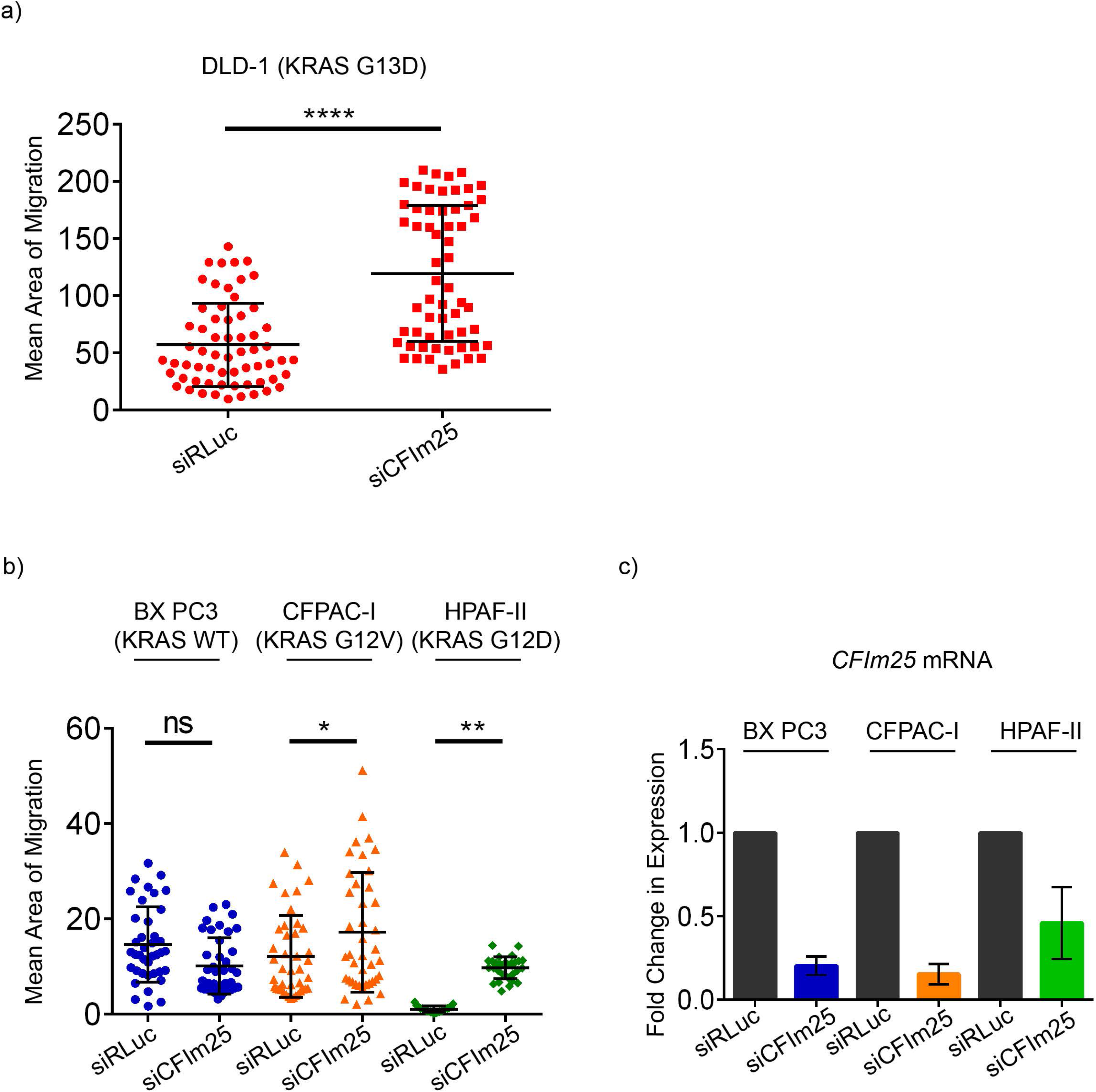
a) Quantification of migration in DLD-1 (KRAS G13D cells). Each data point represents quantification of mean area acquired from at least two independent experiments. b) Quantification of migration in pancreatic cancer cell lines. Each data point represents quantification of mean area acquired from two independent experiments. c) Validation of CFIm25 knockdown by qPCR. n=2 ±SD ****p<0.0001, **p<0.01, *p<0.05, ns=not significant, One-way ANOVA with Tukey’s post hoc test

**Fig. 7.**
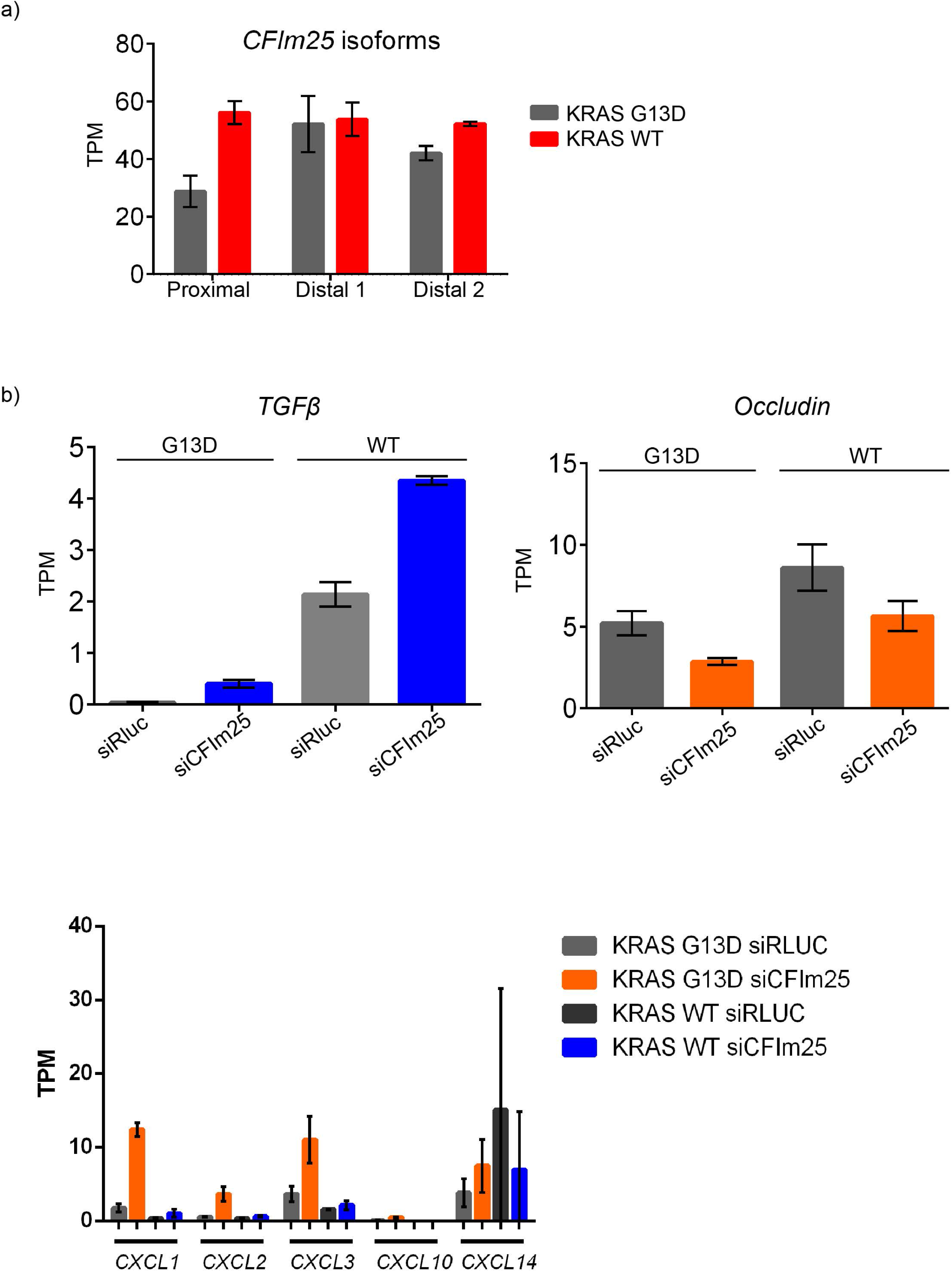
a) KRAS WT cells show higher expression of proximal isoform of CFIm25 as compared to KRAS G13D cells. b) Mining of RNAseq data shows upregulation of *TGβ2*, decreased expression of *Occludin* as well as increase in expression of EMT associated chemokines. TPM represents transcripts per million, n=3 ±SD

**Fig. 8.**
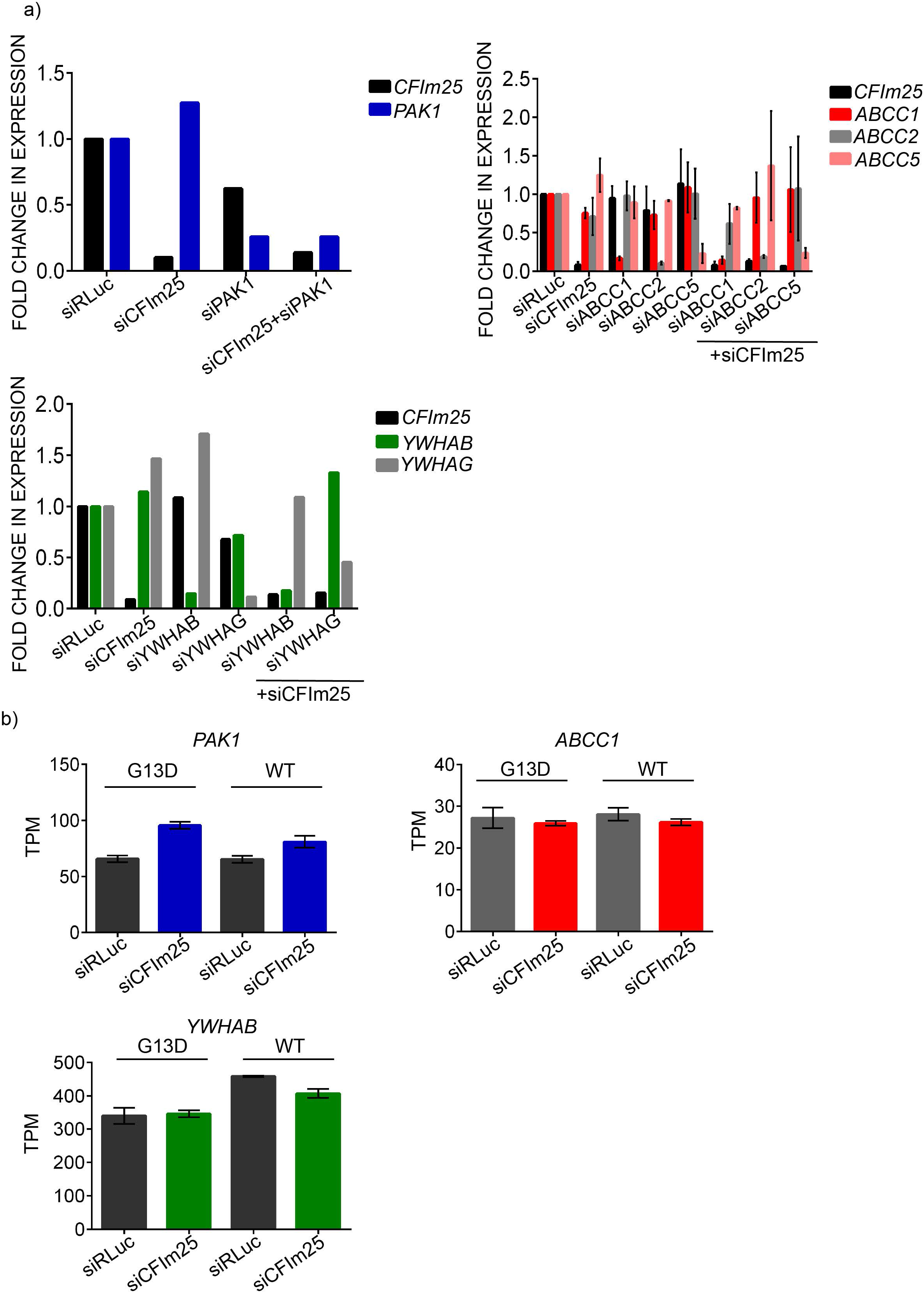
a) Representative qPCR quantification validating knockdown of CFIm25 and/or indicated APA-regulated transcript normalized to *18S* as housekeeping control. b) Analysis of overall gene expression for *PAK1, ABCC1* and *YWHAB*. TPM represents transcripts per million (TPM), n=3 ±SD

## Notes

### Competing Interest Statement

The authors have declared no competing interest.

## REFERENCES

1. Sud, A., Kinnersley, B. & Houlston, R. S. Genome-wide association studies of cancer: Current insights and future perspectives. Nature Reviews Cancer 17, 692–704 (2017).

2. Vogelstein, B. et al. Cancer genome landscapes. Science 340, 1546–1558 (2013).

3. Pylayeva-Gupta, Y., Grabocka, E. & Bar-Sagi, D. RAS oncogenes: Weaving a tumorigenic web. Nature Reviews Cancer 11, 761–774 (2011).

4. Haigis, K. Differential effects of oncogenic K-Ras and N-Ras on proliferation, differentiation and tumor progression in the colon. Nat. Genet. 40, 600–608 (2008).

5. Hobbs, G. A., Der, C. J. & Rossman, K. L. RAS isoforms and mutations in cancer at a glance. J. Cell Sci. 129, 1287–1292 (2016).

6. Tuveson, D. Endogenous oncogenic K-ras(G12D) stimulates proliferation and widespread neoplastic and developmental defects. Cancer Cell 5, 375–387 (2004).

7. Fragoso, R. et al. Modulating the Strength and Threshold of NOTCH Oncogenic Signals by mir-181a-1/b-1. PLoS Genet. 8, (2012).

8. Murphy, D. J. et al. Distinct Thresholds Govern Myc’s Biological Output In Vivo. Cancer Cell 14, 447–457 (2008).

9. Giam, M. & Rancati, G. Aneuploidy and chromosomal instability in cancer: A jackpot to chaos. Cell Division 10, (2015).

10. Ben-David, U. & Amon, A. Context is everything: aneuploidy in cancer. Nature Reviews Genetics 21, 44–62 (2020).

11. Sternberg, P. W. & Han, M. Genetics of RAS signaling in C. elegans. Trends in Genetics 14, 466–472 (1998).

12. Tax, F. E., Thomas, J. H., Ferguson, E. L. & Horvitz, H. R. Identification and characterization of genes that interact with lin-12 in Caenorhabditis elegans. Genetics 147, 1675–95 (1997).

13. Han, M., Aorian, R. V. & Sternberg, P. W. The let-60 locus controls the switch between vulval and nonvulval cell fates in Caenorhabditis elegans. Genetics 126, 899–913 (1990).

14. Hajnal, A., Whitfield, C. W. & Kim, S. K. Inhibition of Caenorhabditis elegans vulval induction by gap-1 and by let-23 receptor tyrosine kinase. Genes Dev. 11, 2715–2728 (1997).

15. Jongeward, G. D., Clandinin, T. R. & Sternberg, P. W. sli-1, a negative regulator of let-23-mediated signaling in C. elegans. Genetics 139, 1553–66 (1995).

16. Clark, S. G., Stern, M. J. & Horvritz, H. R. C. elegans cell-signalling gene sem-5 encodes a protein with SH2 and SH3 domains. Nature 356, 340–344 (1992).

17. Cox, A. D., Fesik, S. W., Kimmelman, A. C., Luo, J. & Der, C. J. Drugging the undruggable RAS: Mission Possible? Nature Reviews Drug Discovery 13, 828–851 (2014).

18. Johnston, S. R. D. Farnesyl transferase inhibitors: A novel targeted therapy for cancer. Lancet Oncology 2, 18–26 (2001).

19. Ryan, M. B. & Corcoran, R. B. Therapeutic strategies to target RAS-mutant cancers. Nature Reviews Clinical Oncology 15, 709–720 (2018).

20. Kirkwood, J. M. et al. Phase II, open-label, randomized trial of the MEK1/2 inhibitor selumetinib as monotherapy versus temozolomide in patients with advanced melanoma. Clin. Cancer Res. 18, 555–567 (2012).

21. LoRusso, P. M. et al. Phase I pharmacokinetic and pharmacodynamic study of the oral MAPK/ERK kinase inhibitor PD-0325901 in patients with advanced cancers. Clin. Cancer Res. 16, 1924–1937 (2010).

22. Flaherty, K. T. et al. Improved survival with MEK inhibition in BRAF-mutated melanoma. N. Engl. J. Med. 367, 107–114 (2012).

23. Downward, J. Targeting RAS signalling pathways in cancer therapy. Nature Reviews Cancer 3, 11–22 (2003).

24. Sun, C. et al. Intrinsic Resistance to MEK Inhibition in KRAS Mutant Lung and Colon Cancer through Transcriptional Induction of ERBB3. CellReports 7, 86–93 (2014).

25. Van Allen, E. M. et al. The genetic landscape of clinical resistance to RAF inhibition in metastatic melanoma on behalf of DeCOG. Cancer Discov 4, 94–109 (2014).

26. Di Giammartino, D. C., Nishida, K. & Manley, J. L. Mechanisms and Consequences of Alternative Polyadenylation. Molecular Cell 43, 853–866 (2011).

27. Mayr, C. What are 3′ utrs doing? Cold Spring Harb. Perspect. Biol. 11, a034728 (2019).

28. Berkovits, B. D. & Mayr, C. Alternative 3′ UTRs act as scaffolds to regulate membrane protein localization. Nature 522, 363–367 (2015).

29. Sandberg, R., Neilson, J. R., Sarma, A., Sharp, P. A. & Burge, C. B. Proliferating cells express mRNAs with shortened 3′ untranslated regions and fewer microRNA target sites. Science (80-.). 320, 1643–1647 (2008).

30. Jan, C. H., Friedman, R. C., Ruby, J. G. & Bartel, D. P. Formation, regulation and evolution of Caenorhabditis elegans 3’UTRs. Nature 469, 97–103 (2011).

31. Derti, A. et al. A quantitative atlas of polyadenylation in five mammals. Genome Res. 22, 1173–1183 (2012).

32. Mayr, C. & Bartel, D. P. Widespread Shortening of 3′UTRs by Alternative Cleavage and Polyadenylation Activates Oncogenes in Cancer Cells. Cell 138, 673–684 (2009).

33. Xia, Z. et al. Dynamic analyses of alternative polyadenylation from RNA-seq reveal a 3’2-UTR landscape across seven tumour types. Nat. Commun. 5, 5274 (2014).

34. Elkon, R. et al. E2F mediates enhanced alternative polyadenylation in proliferation. Genome Biol. 13, R59 (2012).

35. Han, M., Aorian, R. V. & Sternberg, P. W. The let-60 locus controls the switch between vulval and nonvulval cell fates in Caenorhabditis elegans. Genetics 126, 899–913 (1990).

36. Perrin, A. J. et al. Noncanonical control of C. elegans germline apoptosis by the insulin-IGF-1 and Ras-MAPK signaling pathways. Cell Death Differ. 20, 97–107 (2013).

37. Battu, G., Hoier, E. F. & Hajnal, A. The C. elegans G-protein-coupled receptor SRA-13 inhibits RAS/MAPK signalling during olfaction and vulval development. Development 130, 2567–2577 (2003).

38. Honnen, S. J. et al. C. elegans VANG-1 modulates life span via insulin/IGF-1-Like signaling. PLoS One 7, (2012).

39. Schuster, E. et al. DamID in C. elegans reveals longevity-associated targets of DAF-16/FoxO. Mol. Syst. Biol. 6, 399 (2010).

40. Ookuma, S., Fukuda, M. & Nishida, E. Identification of a DAF-16 transcriptional target gene, scl-1, that regulates longevity and stress resistance in Caenorhabditis elegans. Curr. Biol. 13, 427–431 (2003).

41. Murphy, C. T. et al. Genes that act downstream of DAF-16 to influence the lifespan of Caenorhabditis elegans. Nature 424, 277–284 (2003).

42. Seung, W. O. et al. Identification of direct DAF-16 targets controlling longevity, metabolism and diapause by chromatin immunoprecipitation. Nat. Genet. 38, 251–257 (2006).

43. Pinkston-Gosse, J. & Kenyon, C. DAF-16/FOXO targets genes that regulate tumor growth in Caenorhabditis elegans. Nat. Genet. 39, 1403–1409 (2007).

44. McElwee, J., Bubb, K. & Thomas, J. H. Transcriptional outputs of the Caenorhabditis elegans forkhead protein DAF-16. Aging Cell 2, 111–121 (2003).

45. Kubo, T., Wada, T., Yamaguchi, Y., Shimizu, A. & Handa, H. Knock-down of 25 kDa subunit of cleavage factor Im in Hela cells alters alternative polyadenylation within 3’-UTRs. Nucleic Acids Res. 34, 6264–6271 (2006).

46. Lianoglou, S., Garg, V., Yang, J. L., Leslie, C. S. & Mayr, C. Ubiquitously transcribed genes use alternative polyadenylation to achieve tissue-specific expression. Genes Dev. 27, 2380–2396 (2013).

47. Shirasawa, S., Furuse, M., Yokoyama, N. & Sasazuki, T. Altered growth of human colon cancer cell lines disrupted at activated Ki-ras. Science (80-.). 260, 85–88 (1993).

48. Deer, E. L. et al. Phenotype and genotype of pancreatic cancer cell lines. Pancreas 39, 425–435 (2010).

49. Ha, K. C. H., Blencowe, B. J. & Morris, Q. QAPA: A new method for the systematic analysis of alternative polyadenylation from RNA-seq data. Genome Biol. 19, (2018).

50. Wang, N. et al. CXCL1 derived from tumor-associated macrophages promotes breast cancer metastasis via activating NF-κB/SOX4 signaling. Cell Death Dis. 9, 880 (2018).

51. Ikenouchi, J., Matsuda, M., Furuse, M. & Tsukita, S. Regulation of tight junctions during the epithelium-mesenchyme transition: Direct repression of the gene expression of claudins/occludin by Snail. J. Cell Sci. 116, 1959–1967 (2003).

52. Sarvaiya, P. J., Guo, D., Ulasov, I., Gabikian, P. & Lesniak, M. S. Chemokines in tumor progression and metastasis. Oncotarget 4, 2171–2185 (2013).

53. Kim, W., Underwood, R. S., Greenwald, I. & Shaye, D. D. Ortholist 2: A new comparative genomic analysis of human and caenorhabditis elegans genes. Genetics 210, 445–461 (2018).

54. Wernicke, C. M. et al. MondoA is highly overexpressed in acute lymphoblastic leukemia cells and modulates their metabolism, differentiation and survival. Leuk. Res. 36, 1185–1192 (2012).

55. He, D. X. et al. Methylation-regulated miR-149 modulates chemoresistance by targeting GlcNAc N-deacetylase/N-sulfotransferase-1 in human breast cancer. FEBS J. 281, 4718–4730 (2014).

56. Tivnan, A. et al. Inhibition of multidrug resistance protein 1 (MRP1) improves chemotherapy drug response in primary and recurrent glioblastoma multiforme. Front. Neurosci. 9, (2015).

57. Parsons, B. L., McKim, K. L. & Myers, M. B. Variation in organ-specific *PIK3CA* and *KRAS* mutant levels in normal human tissues correlates with mutation prevalence in corresponding carcinomas. Environ. Mol. Mutagen. 58, 466–476 (2017).

58. Mueller, S. et al. Evolutionary routes and KRAS dosage define pancreatic cancer phenotypes. Nature 554, 62–68 (2018).

59. Anglesio, M. S. et al. Cancer-Associated Mutations in Endometriosis without Cancer. N. Engl. J. Med. 376, 1835–1848 (2017).

60. Masamha, C. P. et al. CFIm25 links alternative polyadenylation to glioblastoma tumour suppression. Nature 510, 412–416 (2014).

61. Wang, Y. et al. CFIm25 inhibits hepatocellular carcinoma metastasis by suppressing the p38 and JNK/c-Jun signaling pathways. Oncotarget 9, 11783–11793 (2018).

62. Tan, S. et al. NUDT21 negatively regulates PSMB2 and CXXC5 by alternative polyadenylation and contributes to hepatocellular carcinoma suppression. Oncogene 37, 4887–4900 (2018).

63. Lou, J. C. et al. Silencing NUDT21 attenuates the mesenchymal identity of glioblastoma cells via the NF-κB pathway. Front. Mol. Neurosci. 10, (2017).

64. Chu, Y. et al. Nudt21 regulates the alternative polyadenylation of Pak1 and is predictive in the prognosis of glioblastoma patients. Oncogene 38, 4154–4168 (2019).

65. Lo, H.-W. Targeting Ras-RAF-ERK and its Interactive Pathways as a Novel Therapy for Malignant Gliomas. Curr. Cancer Drug Targets 10, 840 (2010).

66. Cole, S. P. C. et al. Pharmacological Characterization of Multidrug Resistant MRP-transfected Human Tumor Cells. Cancer Res. 54, 5902–5910 (1994).

67. Berger, W. et al. Multidrug resistance markers P-glycoprotein, multidrug resistance protein 1, and lung resistance protein in non-small cell lung cancer: Prognostic implications. J. Cancer Res. Clin. Oncol. 131, 355–363 (2005).

68. Nooter, K. et al. The prognostic significance of expression of the multidrug resistance-associated protein (MRP) in primary breast cancer. Br. J. Cancer 76, 486–493 (1997).

69. Henderson, M. J. et al. ABCC multidrug transporters in childhood neuroblastoma: Clinical and biological effects independent of cytotoxic drug efflux. J. Natl. Cancer Inst. 103, 1236–1251 (2011).

70. Yamada, A. et al. ABCC1-exported sphingosine-1-phosphate, produced by sphingosine kinase 1, shortens survival of mice and patients with breast cancer. Mol. Cancer Res. 16, 1059–1070 (2018).

71. Gao, M. et al. MiR-145 sensitizes breast cancer to doxorubicin by targeting multidrug resistance-associated protein-1. Oncotarget 7, 59714–59726 (2016).

72. Lee, S. H. et al. Widespread intronic polyadenylation inactivates tumour suppressor genes in leukaemia. Nature 561, 127–131 (2018).

73. ten Berge, D., Brugmann, S. A., Helms, J. A. & Nusse, R. Wnt and FGF signals interact to coordinate growth with cell fate specification during limb development. Development 135, 3247–3257 (2008).

74. Sun, Y. et al. Treatment-induced damage to the tumor microenvironment promotes prostate cancer therapy resistance through WNT16B. Nat. Med. 18, 1359–1368 (2012).

75. Pommier, Y. Topoisomerase I inhibitors: Camptothecins and beyond. in Nature Reviews Cancer 6, 789–802 (Nature Publishing Group, 2006).

76. Ma, B. et al. Reciprocal regulation of integrin β4 and KLF4 promotes gliomagenesis through maintaining cancer stem cell traits. J. Exp. Clin. Cancer Res. 38, 23 (2019).

77. Bierie, B. et al. Integrin-β4 identifies cancer stem cell-enriched populations of partially mesenchymal carcinoma cells. Proc. Natl. Acad. Sci. U. S. A. 114, E2337–E2346 (2017).

78. Kiran, S., Dar, A., Singh, S. K., Lee, K. Y. & Dutta, A. The Deubiquitinase USP46 Is Essential for Proliferation and Tumor Growth of HPV-Transformed Cancers. Mol. Cell 72, 823–835.e5 (2018).

79. Li, X. et al. The deubiquitination enzyme USP46 functions as a tumor suppressor by controlling PHLPP-dependent attenuation of Akt signaling in colon cancer. Oncogene 32, 471–478 (2013).

80. Wagner, E. F. & Nebreda, Á. R. Signal integration by JNK and p38 MAPK pathways in cancer development. Nature Reviews Cancer 9, 537–549 (2009).

81. De Sury, R., Martinez, P., Procaccio, V., Lunardi, J. & Issartel, J. P. Genomic structure of the human NDUFS8 gene coding for the iron-sulfur TYKY subunit of the mitochondrial NADH:ubiquinone oxidoreductase. Gene 215, 1–10 (1998).

82. Luo, D. et al. PPA1 promotes NSCLC progression via a JNK- and TP53-dependent manner. Oncogenesis 8, 1–13 (2019).

83. Yang, Y. et al. Inorganic pyrophosphatase (PPA1) is a negative prognostic marker for human gastric cancer. Int. J. Clin. Exp. Pathol. 8, 12482–12490 (2015).

84. Subramani, D. & Alahari, S. K. Integrin-mediated function of Rab GTPases in cancer progression. Molecular Cancer 9, 312 (2010).

85. Tari, A. M., Hung, M. C., Li, K. & Lopez-Berestein, G. Growth inhibition of breast cancer cells by Grb2 downregulation is correlated with inactivation of mitogen-activated protein kinase in EGFR, but not in ErbB2, cells. Oncogene 18, 1325–1332 (1999).

86. Gabrielli, F., Donadel, G., Bensi, G., Heguy, A. & Melli, M. A Nuclear Protein, Synthesized in Growth-Arrested Human Hepatoblastoma Cells, is a Novel Member of the Short-Chain Alcohol Dehydrogenase Family. Eur. J. Biochem. 232, 473–477 (1995).

87. Radu, M., Semenova, G., Kosoff, R. & Chernoff, J. PAK signalling during the development and progression of cancer. Nature Reviews Cancer 14, 13–25 (2014).

88. Chow, H. Y. et al. Group I Paks are essential for epithelial-mesenchymal transition in an Apc-driven model of colorectal cancer. Nat. Commun. 9, (2018).

89. Lee, J., Ogushi, S., Saitou, M. & Hirano, T. Condensins I and II are essential for construction of bivalent chromosomes in mouse oocytes. Mol. Biol. Cell 22, 3465–3477 (2011).

90. Dickinson, D. J., Pani, A. M., Heppert, J. K., Higgins, C. D. & Goldstein, B. Streamlined genome engineering with a self-excising drug selection cassette. Genetics 200, 1035–1049 (2015).

91. Collado-Torres, L. et al. Flexible expressed region analysis for RNA-seq with derfinder. Nucleic Acids Res. 45, 9 (2016).

92. Anders, S., Reyes, A. & Huber, W. Detecting differential usage of exons from RNA-seq data. Genome Res. 22, 2008–2017 (2012).

93. Love, M. I., Huber, W. & Anders, S. Moderated estimation of fold change and dispersion for RNA-seq data with DESeq2. Genome Biol. 15, 550 (2014).

94. Ha, K. C. H., Blencowe, B. J. & Morris, Q. QAPA: a new method for the systematic analysis of alternative polyadenylation from RNA-seq data. Genome Biol. 19, 45 (2018).

95. Harrow, J. et al. GENCODE: The reference human genome annotation for the ENCODE project. Genome Res. 22, 1760–1774 (2012).

96. Patro, R., Mount, S. M. & Kingsford, C. Sailfish enables alignment-free isoform quantification from RNA-seq reads using lightweight algorithms. Nat. Biotechnol. 32, 462–464 (2014).

